# p53 status determines the epigenetic response to demethylating agents Azacitidine and Decitabine

**DOI:** 10.1101/2025.05.19.654807

**Authors:** Emma Langdale Hands, Arndt Wallmann, Gabrielle Oxley, Sophie Storrar, Rochelle D’Souza, Mathew Van de Pette

## Abstract

5’-Azacitidine (Aza) and 5-Aza-2’-deoxycytidine (Dac) are widely used demethylating drugs that directly integrate into nucleic acids. They are frequently used interchangeably, surprisingly as their selectivity is unique from the other, with no predictors of response or clinical biomarkers to indicate drug preference. Using these drugs to induce demethylation, we combine DRIPc-Seq, Immunostaining, RNA-Seq and Mass spectrometry to uncover unique cellular responses. Activation of p53, exclusively by Aza, sustained accumulation of R-loops in CpG islands of *p53* target genes. This effect was abolished by the removal of *p53*, compounded by destabilisation of heterochromatin marks. Dac treatment induced global chromatin modification, sustaining DNA damage, which was heightened in the absence of p53. Rescue experiments reversed the changes observed in the epigenome, demonstrating a direct role for p53 in preserving H3K9me3 and H3K27me3. These insights further our knowledge of how cells recognize and respond to methylation changes and uncover novel roles for p53 in modulation of the epigenome. Further to this, we determine a first in kind biomarker in p53 status that may be relevant for clinical settings.

## Introduction

The scope of our ability to target epigenetic modifications through pharmaceutical agents has expanded significantly in recent years, including Polycomb Repressor Complex II inhibitors, histone deacetylase inhibitors and drugs that specifically modify RNA methylation status (1, 2). Many of these show great promise, however it is only DNA methyltransferase inhibitors that have been progressed into the clinic to date. 5’-Azacitidine (Aza) and 5-Aza-2’-deoxycytidine (Dac) have been used for the treatment of Acute Myeloid Leukaemia (AML) and Myelodysplastic Syndrome (MDS) for a number of years (3–5) and are currently being tested for their efficacy in solid tumour treatment, often in combination with other therapies (6, 7). Their mode of action differs, by virtue of the ability of Aza to integrate into RNA and DNA, inhibiting m5C methylation in both, whereas Dac exclusively integrates into DNA (8, 9). While their ability to inhibit tumorigenesis is well-known, the cellular pathways through which this is achieved remains opaque, beyond their ability to reduce the hypermethylation that is generally associated with the cancer state (10). This is troubling as these drugs are frequently used interchangeably in the clinic, and there is no known predictive marker that would indicate which drug to prescribe to ensure maximal efficacy (11, 12).

Studies have indicated that in addition to a global removal of DNA methylation, Aza and Dac also induce broader changes to the epigenome following treatment (13–16). Most recently, treatment of a cancer cell line with Dac induced an abundance of R-loops (17). R-loops are transient and unstable tripartite complexes that form primarily when a nascent RNA hybridises with one of the DNA strands, ejecting the other. This structure carries with it risks of DNA damage through double strand breaks, as transcriptional stalling caused by the R-loop can lead to a collision of the replication and transcription complexes (18, 19), thereby making any cancer treatment that induces an excess of R-loops to be potentially counterproductive. However, the relative abundance and distribution of R-loops in otherwise healthy cells, observed through genome-wide sequencing techniques (20, 21), demonstrates that with the risk must come the reward of such a structure. Indeed, multiple studies in recent years have revealed physiological roles of R-loops across different genes and cell types (22–26).

The physiological function of the bulk of R-loops are thought to partly converge on transcriptional regulation, whether that be through modifying the total dosage of a gene product, or by altering the splicing pattern of the transcribed RNAs; the manner in which an R-loop achieves this functional consequence is believed to be contingent on its location within any given gene locus (21, 27, 28). In general, so-called promoter R-loops are considered to be transcriptionally activating (23), while terminator R-loops are shown to induce repressive chromatin marks and are associated with DNA damage inducing events (28, 29). In the literature, R-loops are frequently characterised as being either physiological, and required for normal cell function, or pathological, forming aberrantly and with the potential to induce DNA damage. However there is currently little consensus as to what constitutes, or how to distinguish, physiological or pathological R-loops. All R-loops will need to be removed however and to achieve this, cellular machinery primarily mediated by RNAse H (30, 31), resolves them, resulting in a highly dynamic mark.

The variation on the functional consequence of an R-loop depending on where it forms, and the overriding modification of transcription, independent of changes to genetic sequence, implicate R-loops as a component of the overall epigenetic architecture of a cell. Further to this, emerging data has highlighted that R-loop formation and removal is responsive to, and potentially responsible for, changes to distinct epigenetic marks, including DNA methylation and histone modifications (28, 32, 33). In particular, R-loops are especially enriched in CpG islands that are unmethylated, with recent data suggesting that changes in methylation will consequently modify the global R-loop profile (21, 34). With this in mind, we asked whether depletion of the methylome, through either Aza or Dac, would consequently involve the genome-wide accumulation of R-loops in newly unmethylated CpG islands, and whether this results in elevated DNA damage and cell stress. Recent advances in methodology now allows for strand specific, single-base resolution mapping of R-loops genome-wide, through the next-generation sequencing (NGS) of the offending RNA strand (DNA-RNA immunoprecipitation followed by cDNA conversion and sequencing, DRIPc-Seq (20)). Here we combine this approach with epigenetic analysis to uncover divergent mechanisms of cellular action for Aza and Dac.

## Methods

### Cell Culture

The Human embryonic kidney cell line (HEK293) and a HEK293 p53 KO cell line were used for the majority of experiments in this study. Both cell lines were cultured in Dulbecco’s Modified Eagle Medium (DMEM, Gibco) supplemented with 1% glutamax (Gibco), 10% fetal bovine serum (FBS, Gibco) and 1% Penicillin and Streptomycin (Gibco). Cells were grown under standard conditions at 37℃ and 5% CO_2_. Cells were passaged every 2-3 days at a 1:10 dilution.

A p53-inducible H1299 cell line, from here on called ip53 H1299, was kindly gifted by Masashi Narita’s lab (Cancer Research UK Cambridge Institute). Information on the establishment of the cell line can be found in (35). ip53 H1299 cells were cultured in growth medium (DMEM supplemented with 10% FBS) under standard conditions at 37℃ and 5% CO_2_. To induce the expression of wt p53 in these cells, cells were seeded into growth media containing 100ng/mL Doxycycline (Dox) in DMSO, with cells not receiving Dox being treated with the same volume of DMSO, as a solvent control. Dox treatment was refreshed every 24hrs as necessary for the desired length of treatment.

### Drug Treatment

The drugs 5’-Azacitidine (Azacitidine) (Sigma, A2385) and 5-aza-2’-deoxycitidine (Decitabine) (Cell Guidance systems, SM46-5) were dissolved in acetic acid and water (1:1) and stored as 5mg/mL stocks at -80°C for a maximum of one month. HEK293, HEK293 p53 KO cells or ip53 H1299 cells were plated at 70% confluence and left for 24hrs before being treated with 1.5µM Azacitidine, Decitabine or solvent for 48hrs, with the treatment refreshed after 24hrs. The exception to this being for the time course experiments where these cells were treated with either drug for the stated amount of time and the toxicity study where these cells were treated with a range of doses of either drug.

### RNA Extraction

RNA was extracted from cells using the RNeasy Mini Plus Kit (QIAGEN, #24136) following kit instructions with the additional homogenisation step using the QIAshredder (QIAGEN, #79656), for the majority of experiments. RNA was quantified on a NanoDrop spectrometer or Qubit (Thermo Fisher) with the HS RNA Kit. RNA was extracted for the qPCR Time Course Experiment using Trizol. 1mL of Trizol was added directly to cells and incubated at room temperature for 5 minutes. Cell lysate was transferred into an 1.5ml tube and 0.2mL of 100% chloroform added, vortexed well and incubated at room temperature for 2-3 minutes. Samples were then centrifuged at no more than 12, 000*g* for 15 minutes at 2-8℃. The aqueous phase was retained and washed with 0.5mL of 100% isopropanol at room temperature for 10 minutes. Samples were then centrifuged at 12, 000*g* for 10 minutes at 2-4℃ to pellet the RNA. The RNA pellet was then washed twice in 1mL of 75% ethanol before being allowed to dry for 5-10 minutes. RNA was then resuspended in nuclease-free water and stored in -80℃.

### qPCR

RNA was converted to cDNA using the ReverAid RT Reverse Transcription Kit (Thermo Fisher, K1691) as per kit instructions with an input of 2µg of RNA. qPCR reactions were performed with 2µL of cDNA, oligonucleotides targeting specific genes, prepared as 10µM working solutions, and Fast SYBR^TM^ Green Master Mix (Thermo Fisher). Reactions were prepared as per the protocol from Thermo Fisher for the Fast SYBR Green Master Mix. qPCR was performed using the Thermo Fisher QuantStudio 6 Flex under the following cycling conditions: 95℃ 20 sec, 95℃ 3 sec, 60℃ 30 sec; for 40 cycles, followed by a melt curve to validate the primers used. Negative controls included a no template (no cDNA) reaction and minus RT reaction. GAPDH was used as a housekeeping gene for all qPCR experiments. Quantification of qPCR was performed by correcting the CT mean against the housekeeping gene to produce a ΔCT. From this, ΔΔCT and RQ values were calculated compared to the control treatment group.

### Western Blot

Protein was extracted from cells by lysing cells directly with RIPA buffer (ThermoFisher #89900) supplemented with 1X protease inhibitor cocktail (cOmplete Protease Inhibitor cocktail, Sigma) and 1X protein phosphatase inhibitor-2 (Sigma), for 10 mins on ice. To quantify protein concentration a BCA assay was performed using the ThermoFisher Pierce^TM^ Protein Assay Kit as per kit instructions and using known standards.

20µg of protein lysate was prepared with 4X Laemmli sample buffer (Bio-Rad), supplemented with 10% β-mercaptoethanol. Denatured protein was run using NuPage 4-12% Bis-Tris Gels (Thermo Fisher Scientific). Samples were transferred onto a PVDF membrane. Membranes were then blocked with either BSA or 5% Milk in Tris Buffered Saline (TBS) with 1% Tween (TBS-T) for 2 hours at room temperature. Specific proteins were detected by incubating membranes in blocking buffer with primary antibodies (listed in table) at 4℃ for 12-20 hours. Licor IRDye (Infra-red Fluorescent) secondary antibodies at 20, 000X dilution were used for 30 minutes at room temperature. Antibody detection was carried out on a Licor Odessey CLx with signal analysis obtained from Licor ImageStudio software.

### DRIP-Mass Spectrometry

To analyse the R-loop interactome, DNA:RNA immunoprecipitation (DRIP) was coupled with Label Free relative quantification Mass Spectrometry. Samples were obtained for Mass Spectrometry from HEK293 and HEK293 p53 KO cells following treatment with 1.5µM Azacitidine or Decitabine or solvent for 48hrs. Samples were then prepared as stated in (36) with the following changes. Following sonication, one third of each sample was treated with 5.5U of RNaseH per µg of DNA overnight at 37℃. 25µg of the sonicated DNA or RNaseH treated DNA was diluted 1:4 in Resuspension Buffer (RSB) + Tween 20 (RSB+T) buffer. Samples were then subjected to immunoprecipitation with 10µg of S9.6 antibody (or no antibody, as a negative control) and 0.1ng RNase A per µg genomic DNA, and rotated at 4℃ for 4 hours. 50µL protein A/G Dynabeads (Thermo Fisher Scientific) was added to each solution and incubated for a further 2 hours at 4℃ with rotation. Beads were washed four times with RSB+T and then twice with RSB buffer. Samples were then eluted in RapiGest (Fisher Scientific) for 30 minutes at room temperature. Eluted samples were then processed by filter aided sample preparation, and LC-MS/MS, which was kindly performed by the University of Cambridge MRC Toxicology Unit Proteomics facility. In brief, samples were precipitated in a 0.07% β-mercaptoethanol buffer followed by an in-solution digestion with AMBIC and RapiGest, protein reduction in Dithiothreitol (DTT), alkylation with Iodoacetamide and IAA quenched with DTT. Samples were then ready for injection. Samples were injected in a randomised manner.

Data processing was performed by the University of Cambridge MRC Toxicology Unit Proteomics facility. Raw data were imported and data processed in Proteome Discoverer v2.5 (Thermo Fisher Scientific). The raw files were submitted to a database search using Proteome Discoverer with SequestHF and Inferis re-scoring algorithm against the Homo sapiens database containing human protein sequences from UniProt/Swiss-Prot. Common contaminant proteins (several types of human keratins, BSA and porcine trypsin) were added to the database. The spectra identification was performed with the following parameters: MS accuracy, 10 p.p.m.; MS/MS accuracy of 0.01 Da for spectra acquired in Orbitrap analyzer; up to two missed cleavage sites allowed; carbamidomethylation of cysteine as a fixed modification; and oxidation of methionine as variable modifications. Percolator node was used for false discovery rate estimation and only rank 1 peptide identifications of high confidence (FDR<1%) were accepted. PD normalisation was performed using all peptide abundances for the normalisation of the samples. Significantly enriched or depleted proteins in the treated samples, identified compared to control treated samples and not present in negative controls, were analysed for protein localisation, protein domain enrichment and gene ontology enrichment via the STRING pathway network software (v.12) (37). Enriched and depleted protein datasets were also analysed in the clusterMaker2 software (v.2.3.4) for Markov cluster algorithm (MCL) (38) clustering to produce functional cluster maps of each protein network.

### Immunofluorescence

HEK293 and HEK293 p53 KO cells were plated at 70% confluence on sanitised glass cover-slips which had been coated in 0.05mg/mL Poly-D-Lysine. ip53 H1299 cells were plated at 70% confluency on uncoated sanitised glass coverslips. All cells were fixed in 100% ice-cold methanol for 10 minutes on ice. Coverslips were blocked in 3% BSA in PBS, except for S9.6 staining in which coverslips were blocked in 3% BSA in 4xSSC. S9.6, ɑ-tubulin and γ-tubulin primary antibodies were stained at 1:200 dilution, whilst H3K9me3 and H3K27me3 were stained at a 1:1000 dilution. DAPI was used as a counterstain to identify nuclei. All image processing and analysis was performed using Fiji software (version 1.54t). Micronuclei were identified by placing a grid over the image and counting the number of micronuclei in each grid cell. The total number of micronuclei in an image was then divided by the total number of nuclei in the image, as identified using the DAPI count macro script in ImageJ. H3K9me3 and H3K27me3 images were analysed using Macro scripts in which only the nuclear signal is measured. This was done by using the DAPI stain image channel to identify nuclei and create a mask which was then applied to other image channels to measure signal intensity (Integrated Density) and signal area only within the DAPI area. R-loop signal intensity (S9.6) and signal area were calculated in the same way, however the additional measurements of foci area and foci number were also recorded. H3K9me3 and H3K27me3 signal intensity was corrected to DAPI signal intensity by dividing the target signal by the DAPI signal for each nucleus, to produce corrected total nuclear fluorescence (CTNF). This was done to correct for any potential cross over of fluorescence excitation between the target channel secondary antibody (488) and the DAPI channel (460). All scale bars on images were added through the Fiji software. Scales of scale bars are described for each figure in the figure legend.

### WGBS sample prep

DNA was extracted from HEK293 cells which had been treated with 1.5µM of Azacitidine or Decitabine or solvent for 48 hours, using the QIAGEN DNeasy Blood and Tissue Extraction Kit, as per kit instructions. Whole Genome Bisulfite Sequencing (WGBS) conversion and library construction was then performed using Zymo-Seq WGBS Library Kit (Zymo Research) as per kit instructions. The Select-a-size DNA/RNA Clean and Concentrator Magbead Kit (Zymo Research) was used when needed to remove primer dimer contamination from samples. Libraries were visualised before sequencing using the High Sensitivity DNA Kit and Chip on a BioAnalyser (Agilent). All libraries were combined into a single pool and sequenced at the University of Cambridge Biochemistry sequencing facility using the Next-Seq 500 (Illumina), 300-cycles 150bp paired-end reads.

### WGBS data processing and data analysis

FastQC (version 0.11.9) was used to assess the overall quality of the sequenced samples, and TrimGalore (version 0.6.6) was used to trim low-quality bases (quality score lower than 20), adapter sequence, and end-repair bases from the 3ʹ end of reads. Bismark (version 0.22.3) was used for alignment to the human hg19 reference genome, deduplication of reads and methylation calling (39). The methylKit R package (version 1.20.0) was used to test WGBS data for differential methylation for the AZA and DAC exposures compared pairwise to the control group (40). Methylation calls were reported for all nucleotides with a read depth of at least 5. In addition to individual CpG sites, we used the function tileMethylCounts to determine methylated/unmethylated base counts over 1000 bp tiling windows across the genome. The calculateDiffMeth function was used to identify differentially methylated CpGs or 1000 bp tiles. To filter for differentially methylated regions (DMRs), we used the getMethylDiff function with a q-value cut-off of 0.01 and a difference cutoff (absolute value of methylation percentage change) of 25. Referencing to other sequencing data (e.g. DRIPc) was achieved through the GRanges (version 1.46.1) R package by transforming the respective regions into GRanges objects on which e.g. overlap operations could be conducted (subsetByOverlaps) (41). The genomation (version 1.26.0) and ChIPseeker (version 1.30.3) R packages were employed to annotate DMRs to the TxDb hg19 genome (TxDb.Hsapiens.UCSC.hg19.knownGene) (42, 43). The function annotateWithFeatures and annotatePeak were used to annotate the abundance of regions within custom and TxDb genetic features, respectively.

### RNA-Seq library preparation, data processing and data analysis

RNA was extracted, as described above using the kit method. The QIAseq Fast Select - rRNA HMR Kit (Qiagen) was used to prepare RNA sequencing libraries, with 1µg RNA as input, as per kit instructions. To remove primer dimers the Select-a-size DNA/RNA Clean and Concentrator Magbead Kit (Zymo Research) was used, with the desired peak 300bp. Libraries were then pooled at 10nM in 10µL. RNA-sequencing (RNA-Seq) was performed using the Next-Seq 500 (Illumina) for 150 cycles paired-end reads, by the University of Cambridge Biochemistry Department sequencing facility. Quality of data in fastq files was assessed using FastQC and low quality bases (quality score lower than 20) and adapter sequences were removed using TrimGalore. Reads were aligned to the human hg19 reference genome using Hisat2 (version 2.2) and raw read counts were generated using featureCounts. The data was normalised for composition bias using the calcNormFactors function from the edgeR R package (version 3.36.0), and normalised again and transformed to log2 counts per million reads (CPM) using the voom function from the limma R package (version 3.50.3) (44, 45). To determine differential expression, the contrast.fit and eBayes limma functions were used to compute estimated coefficients and standard errors, and apply empirical Bayes moderation, respectively. The p-values were adjusted by using the Benjamini-Hochberg false discovery rate (FDR) approach applying an FDR < 0.05 and Log2(fold change) > 1 to determine significance.

### DRIPc-Seq sample prep

DNA:RNA immunoprecipitation with cDNA conversion sequencing (DRIPc-Seq) was performed on HEK293 and HEK293 p53 KO cells, treated with Azacitidine, Decitabine or solvent, as described above. DRIPc-Seq was performed according to the published method (20), with the exception that the genomic DNA was fragmented using sonication to achieve a fragment size of around 300bp. An RNAseH treated control was used as the negative control. Libraries were then prepared using the TruSeq RNA v2 kit (Illumina) by the University of Cambridge Biochemistry sequencing facility.

### DRIPc-Seq data processing and data analysis

All libraries were sequenced at the University of Cambridge Biochemistry sequencing facility using the Next-Seq 500 (Illumina), 150-cycles single-end reads. As described above, FastQC was used to assess the overall quality of the samples, and TrimGalore was used to trim low-quality bases and adapter sequences. Trimmed reads were mapped to UCSC human hg19 genome using Bowtie2 (version 2.4.2) and the mapped reads were deduplicated using the MarkDuplicates command in Picard tools.

Strand-specific R-loop peak calling was performed using MACS3 (version 3.0.0a6) with the --broad --nomodel --broad-cutoff 0.2 --max-gap 350 --extsize 150 settings. The resulting peaks were merged into union R-loop peaks in a strand-specific manner, collapsing the peaks into non-redundant and distinct peaks. We then compiled a high confidence peak list that contained only peaks that are present in at least two datasets and had not remained in any of the RNaseH-treated samples. To identify R-loop signals that emerge upon treatment, the differentially up- or downregulated R-loop signals for all high confidence peaks were determined using the R package DESeq2 (version 1.34.0) with a cutoff Benjamini-Hochberg adjusted P-value (adjusted P-value) < 0.05 and a Log2(fold change) > 0.5 to identify gained R-loops or Log2(fold change) < -0.5 for lost R-loops. Importantly, changes were only determined with a control treatment that was measured alongside the respective treated sample to account for any underlying changes in R-loops architecture between experiments. The resulting peaks were annotated to the hg19 genome using CHIPseeker and genomation and pathway-enrichment was performed using the ReactomePA R package (version 1.9.4) (43, 46).

### Statistical Analysis

Various statistical tests were used in this study, specific tests are described in the corresponding figure legends or methods sections. All error bars shown are 士SEM unless otherwise stated in the figure legend. For all experiments the following nomenclature was used: * p < 0.05, ** p < 0.01, *** p < 0.001, **** p < 0.0001. Unless otherwise stated, statistical analysis was performed in R (v.4.3.2).

### Data Availability

All raw fastq.gz reads from sequencing experiments were uploaded to the Sequence Read Archive (SRA) under the following reference series WGBS: GSE243785, DRIPc-Seq: GSE243786, RNA-Seq: GSE243784.

### Primer sequences

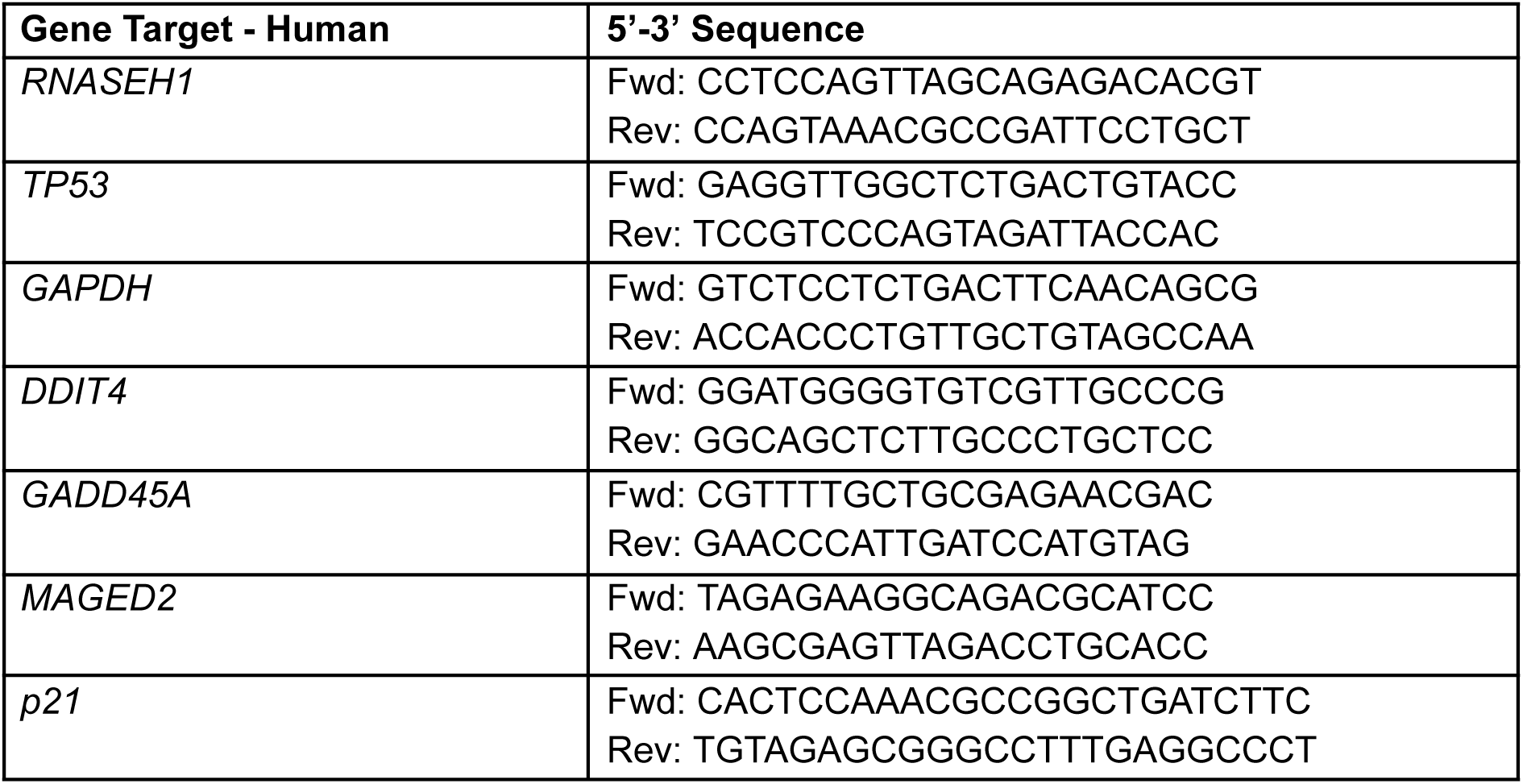

### Antibodies

| Target | Source | Concentration | Blocking Buffer | Use |
| --- | --- | --- | --- | --- |
| H3K9me3 | Abcam ab8898 | 1:1000 | 3% BSA | IF |
| $\beta$ Actin | Abcam ab8337 | 1:1000 | 5% Milk | Western Blot |
| p53 | Abcam ab26 | 1:1000 | 5% BSA | Western Blot |
| Phospho-p53 Ser15 | Cell Signalling #82530 | 1:1000 | 5% BSA | Western Blot |
| S9.6 | MERK MABE1095 | 1:200 | 3% BSA | IF, DRIPc-Seq |
| H3K27me3 | Abcam ab6002 | 1:1000 | 3% BSA | IF |
| $\alpha$ Tubulin | Abcam ab7291 | 1:200 | 3% BSA | IF |
| $\gamma$ Tubulin | Abcam ab11317 | 1:200 | 3% BSA | IF |

| Secondary Antibody | Source | Concentration | Use |
| --- | --- | --- | --- |
| IRDye 680Rd Mouse | Licor 926-68072 | 1:20000 | Western Blot |
| IRDye 680Rd Rabbit | Licor 926-68073 | 1:20000 | Western Blot |
| IRDye 800CW Mouse | Licor 926-32210 | 1:20000 | Western Blot |
| IRDye 800CW Rabbit | Licor 926-32211 | 1:20000 | Western Blot |
| Alexa-Fluor 568 Mouse | ThermoFisher<br>#A-11004 | 1:1000 | IF |
| Alexa-Fluor 388 Rabbit | ThermoFisher<br>#A32731 | 1:1000 | IF |
| Alexa-Fluor 488 Mouse | ThermoFisher<br>#A32723 | 1:1000 | IF |
| DAPI Counterstain | Roche<br>#10236276001 | N/A | IF |

## Results

### Aza and Dac treatment leads to epigenetic re-mapping at sub-toxic concentrations

Aza and Dac are well known to be substantially toxic to mammalian cells, and their anti-cancer properties are thought to occur through the demethylation and activation of tumour suppressor genes (47). While demethylation was a requirement for our model, we sought to identify a clinically relevant dosing regimen for each drug in HEK293 cells where acute toxicity was minimalized (Figure 1A, B). Both drugs were found at 1.5µM to satisfy this requirement, with no significant changes in viability or apoptosis and minimal cytotoxicity after 48hrs of treatment. Whole genome Bisulfite Sequencing (WGBS-Seq) demonstrated that at this dose, substantial genome-wide demethylation occurred following exposure to either drug, with broadly overlapping regions of demethylation. Somewhat surprisingly, given that Aza is believed to preferentially integrate in RNA over DNA (8, 9), this demethylation of the DNA was found to be greater and more widespread in Aza samples than Dac, with 30.6% of significantly demethylated tiles unique to Aza treated cells, compared to only 10.6% in Dac treated cells (Figure 1C). Analysis of the genomic location of all hypomethylated tiles revealed an enrichment in coding regions of the genome, that was predominantly shared by both drugs (Figure 1D, S1A-C).

**Figure 1:**
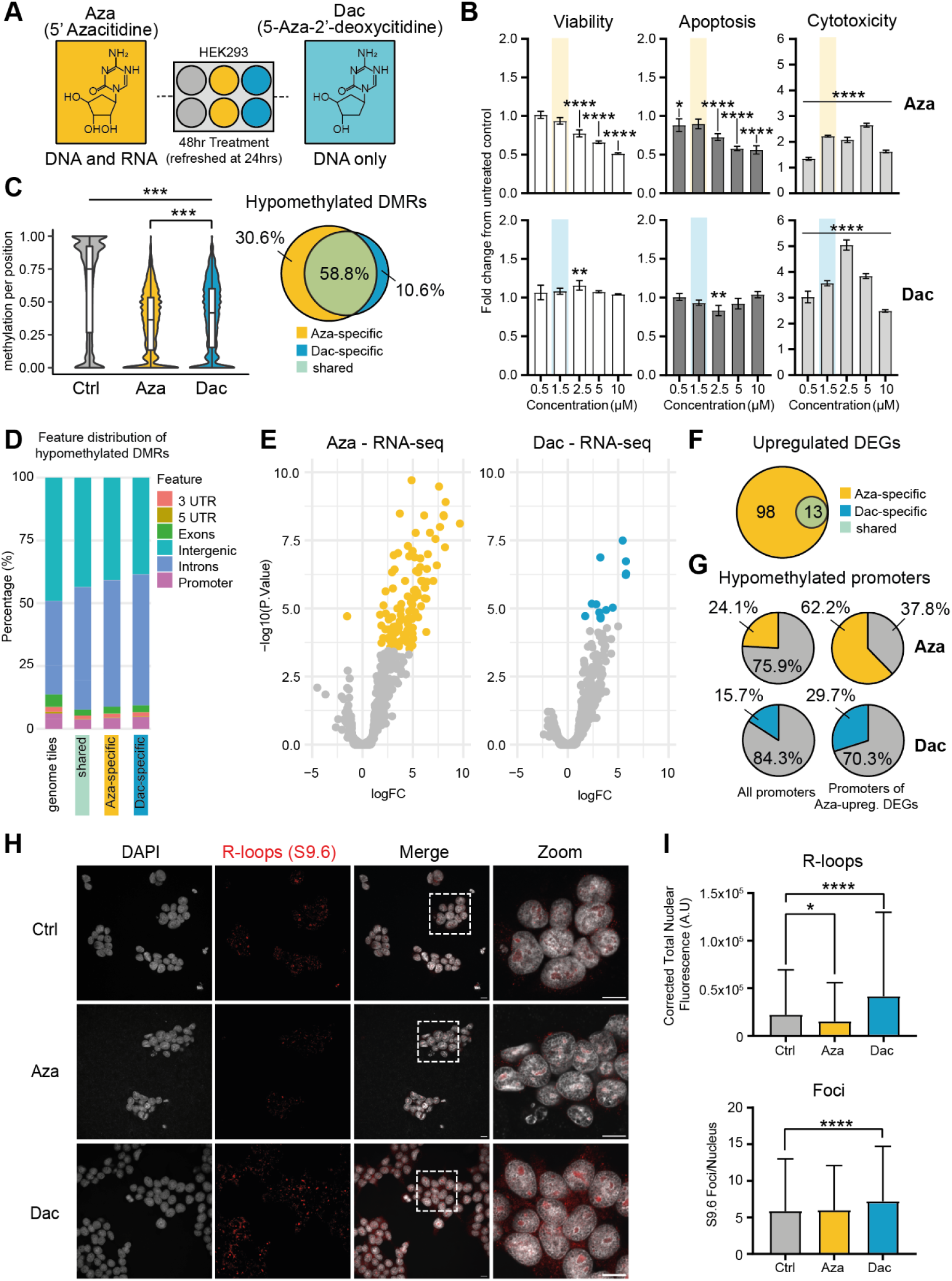
Hypomethylating conditions are sufficient to induce changes in transcription and global R-loop abundance. (**A**) Experimental schematic of drug treatment in HEK293 cells. Cells were treated for 48hr with either Aza, Dac or solvent control with a refresh after 24 hours. (**B**) ApoTox Toxicity assay in HEK293 cells treated with a dose range of Aza or Dac to measure cell viability, apoptosis and cytotoxicity levels. (**C**) Violin plots of CpG methylation of all available positions from WGBS of Aza or Dac treated HEK293 cells. Venn diagram of hypomethylated DMRs based on 1000 bp tiling of the genome. The p values were calculated by one-way ANOVA with Tukey’s correction. ***p < 0.001. (**D**) Feature distribution of all tiles across the genome and shared and treatment-specific DMRs shown in (**C**). (**E**) Volcano-plot of RNA-seq analysis of Aza and Dac treated HEK293 cells. Significantly differentially expressed genes (DEGs) (Aza: 111; Dac: 13) are highlighted in the respective colour. (**F**) Venn diagram of upregulated DEGs in Aza and Dac treated cells. Significantly altered genes are listed in Table S1 (**G**) Proportion of hypomethylated promoters in all genes and the upregulated DEGs under Aza treatment for Aza and Dac treatment, proportion of non-hypomethylated promoters is in grey. (**H**) Representative confocal images using S9.6 antibody to stain Aza and Dac treated HEK293 cells for R-loops (red) and DAPI as a counterstain for DNA (grey). Scale bars 10μm. (**I**) Image quantification of S9.6 staining plotted as corrected total nuclear fluorescence and S9.6 foci number per nuclei. Statistical testing was calculated by one-way ANOVA with Tukey’s correction. *p < 0.05, ****p < 0.0001. Error bars are 士SEM.

To determine if this broad demethylation induced changes in transcription, RNA-Sequencing was performed. 98 genes were uniquely and significantly up-regulated following Aza treatment, in addition to 13 that were also increased following Dac treatment. These 13 genes represented the total of all significantly modified genes in Dac treatments (Figure 1E, F, Table S1). The fact that expression was predominantly elevated genome-wide following the treatment of Aza and Dac corresponds with the known function of methylation and for the 98 genes that were found to be up-regulated following Aza treatment, their promoters were more demethylated when compared to all promoters across the genome (62.2% vs 24.1%). Analysis of these same 98 promoter regions in the Dac treated samples determined that a smaller proportion of these promoters became hypomethylated following that treatment (29.7% vs 15.7%) (Figure 1G). This indicates that beyond the broad demethylation observed by both drugs, the change in methylation status of selected promoters may have a direct impact on the transcriptional profiles of the treated cells.

In light of the known enrichment of R-loops in hypomethylated CpG islands (21, 34), and following our confirmation that Aza and Dac caused global demethylation in the absence of substantial acute toxicity, we asked whether treatment with Aza and Dac induced an accumulation of global R-loops that corresponded to the changes in DNA methylation or transcription. Immunostaining for R-loops (S9.6 clone, red, Figure 1H) demonstrated a differential response to the two drugs, with the total fluorescence and the number of nuclear foci elevated following Dac treatment, in accordance with published work (17), while in contrast a reduction in total fluorescence was observed in Aza treated cells (Figure 1I). This increase in R-loop staining in Dac was accompanied by elevated markers of DNA damage, including micronuclei formation (Figure S1D-E) and mitotic spindle pole errors (Figure S1F-G), which was not observed in Aza treated samples. R-loop formation has been observed to coincide with a decrease in the repressive histone marker H3K27me3 by inhibiting binding of the Polycomb Repressive Complex 2 (PRC2), the canonical writer of H3K27 methylation (48). Immunostaining for H3K27me3 in treated cells showed a significant decrease in nuclear staining intensity in Aza treated cells, whereas there was an increase, though not significant, in Dac treated cells (Figure S1H). In contrast, global H3K9me3 signal was significantly increased in Aza treated cells, while no significant changes were detected in Dac treated cells (Figure S1I).

### Global R-loops are re-mapped following Aza and Dac treatment

Due to the substantially different staining patterns for R-loops in either Aza or Dac treatments, we chose to perform DNA:RNA Immunoprecipitation followed by cDNA conversion and sequencing (DRIPc-Seq) (32). This approach enables single base and strand specific resolution of R-loops, allowing a base resolution genome-wide picture of the changes that had occurred to be generated (Figure 2A). Intra-experimental technical variation has been reported for S9.6 based R-loop detection (Figure S2A) (32, 49) and so to account for this, we employed strict controls as outlined in the methods section. The size and stranded distribution of R-loops were only modestly changed by either treatment (Figure S2B, C). However, the genomic location of Aza or Dac R-loops were substantially disturbed, with the majority of these changes found within coding regions (Figure 2B). These changes though were unique from one another, and while an accumulation of R-loops was found to occur in promoters or at the transcriptional start site (TSS) following Aza treatment, a notable decline in intronic R-loops was also observed. In contrast, promoter R-loops became depleted in Dac samples, while there was a substantial accumulation of R-loops near to the transcriptional termination site (TTS) (Figure 2B, C). In fact, co-accumulation or co-depletion of R-loop peaks in Aza and Dac treatment was almost completely absent, with a much stronger likelihood of an inverse response occurring (Figure S2D).

**Figure 2:**
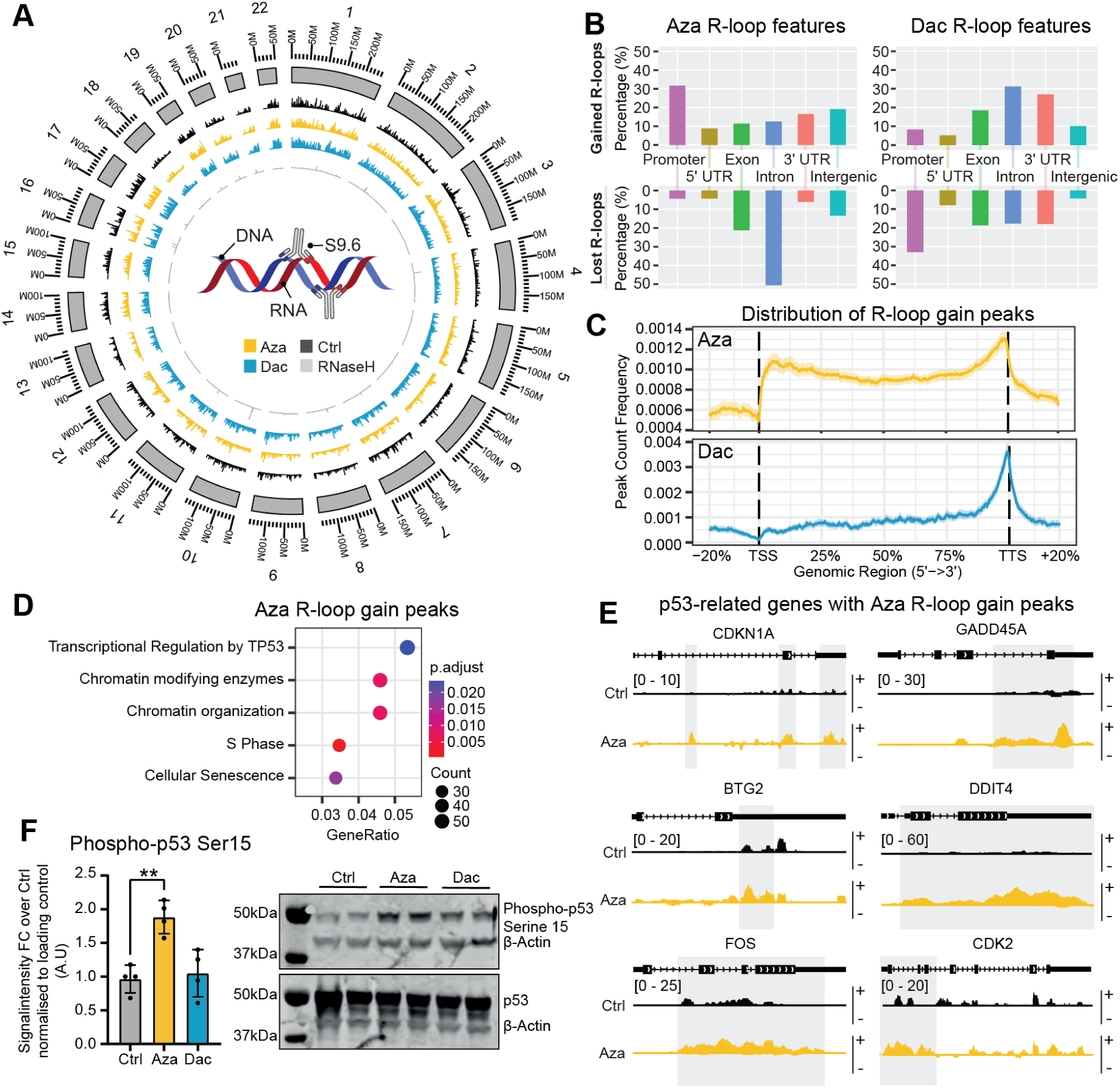
Azacitidine and Decitabine treatment uniquely alter the R-loop landscape. (**A**) Circos plot showing the unstranded R-loop signals across chromosomes in 1000-kb bins for Control (Ctrl), Azacitidine (Aza), Decitabine (Dac) and RNaseH-treated HEK293 cells. Schematic cartoon summarising the DRIPc-seq approach through use of the S9.6 antibody to recognise R-loops which can then be immunoprecipitated for sequencing. (**B**) Feature distribution for significantly increased (gained) and decreased (lost) R-loop signals in cells treated with Aza and Dac. (**C**) Peak count frequency for Aza and Dac gained R-loop peaks across the gene body. (**D**) Top-scoring pathway enrichment terms of genes with gained R-loop peaks from Aza-treated cells. (**E**) DRIPc-seq representative R-loop peak tracks of p53-related genes after Ctrl treatment and Aza treatment (Watson/forward-strand R-loop (+), Crick/reverse-strand R-loop (−)). The gained peaks upon Aza treatment are highlighted. (**F**) Representative Western blot of p53, p53 with serine 15 phosphorylation (Phospho-p53 Ser15) and β-Actin in HEK293 cells treated with 1.5μM Aza or Dac, or solvent control. These images are taken from the same gel. Bar graph of quantification of phospho-p53 Ser15 Western blot signal normalised to total p53 signal, n=4 biological repeats, p values were calculated by one-way ANOVA **p < 0.01, data are presented as mean 土 SEM.

As has been already stated, R-loops are known to preferentially inhabit unmethylated CpG islands (21)(34). In either Aza or Dac treatment however, the relationship between newly demethylated regions and an accumulation of R-loops was not immediately obvious at a genome-wide level. In fact, for Aza treatment, a depletion of an R-loop was more likely to occur in a newly demethylated region, rather than an enrichment, while Dac induced demethylated regions were as likely to form R-loops as they were to lose them (Figure S2E). To overcome this complexity, we restricted our analysis to promoters for Aza treatment, as our earlier analysis had indicated a link between demethylation of promoter regions, and an increase in transcription of the associated gene, that was unique to Aza (Figure 1G). Pathway enrichment for those genes with a newly formed promoter R-loop highlighted genes associated with chromatin architecture and the cell cycle, potentially indicating a global epigenetic response to the initial exposure. To our surprise however, the top enrichment was for p53 target genes (Figure 2D, E). p53 is a protein which is frequently named the ‘guardian of the genome’ because of its various roles in DNA damage repair, cell cycle arrest and apoptosis, in response to a variety of cell stressors, thus making it a critical tumour suppressor and by extension the most frequently mutated gene in human cancers. DNA damage as a result of increased R-loop formation has previously been linked to p53 activation (50), and p53 deficient cells have been shown to be dependant on the DHX9 DexH-box helicase, an RNA helicase that functions in the removal of R-loops (51). Despite this, very little is known about the role of p53 directly in the formation of R-loops.

No p53-target genes had been identified as differentially expressed in the RNA-Seq analysis (Figure 1E, S2F), while p53 is not known to interact directly with R-loops. Previous work has identified that activation of CBP/p300 occurs following Aza treatment (52), and these proteins are known to activate p53 (53). This mechanism would not be predicted to activate p53 following Dac exposure, as it is known to occur through the sensing of RNA modifications and changes to protein synthesis, which Dac is not known to affect. Western blotting for Phospho-p53 Ser 15, a mark that represents activated p53, was performed, and determined to be activated following Aza treatment (Figure 2F). In contrast, such activation was not observed following Dac treatment, and no associated changes in p53-target gene promoter R-loops were detected (Figure 2F, S2G). Promoter R-loops are generally considered to function through the promotion of transcription, however our analysis had not determined transcriptional activation for the p53-target genes in question. By definition, a cis R-loop requires transcription to occur, therefore, we reasoned that it was possible our timing was missing an earlier increase in the expression of these p53-target genes. To determine if this was the cause, we performed time-course analysis of the expression of three known p53-target genes which were enriched for R-loops in the Aza-treated samples (*DDIT4, CDKN1A, GADD45A*) and one p53-target gene which was not enriched for R-loops (*MAGED2*), as well as expression of the p53 gene (*TP53*) (Figure S2H). While modest increases in expression were detected for certain p53-target genes following Aza treatment (*DDIT4, GADD45A*) at timepoints which would have precluded detection through our earlier analysis, a clear concerted increase in other genes (*MAGED2, CDKN1A*) was not observed. Following Dac treatment there was only a significant increase in *MAGED2* expression after 48hrs. Expression of *TP53* was shown to significantly increase following Aza treatment, but not Dac treatment.

### Knockout of p53 affects R-loop formation upon treatment

To determine if p53 had a direct role in regulating R-loop formation following Aza or Dac treatment, R-loop immunofluorescence staining was performed in HEK293 p53 KO cells. Aza and Dac treated p53 KO cells displayed significantly increased total staining and foci number compared to untreated p53 KO cells (Figure 3A, B). The total nuclear staining and foci formation was also significantly higher for both treatments in the KO cells when compared to the WT cells, suggesting that the initial loss of p53 results in its own increase in R-loop formation. Observations of markers of genomic instability were also significantly increased for both treatments in the p53 KO cells compared to WT cells, with a significant increase in micronuclei and abnormal mitotic events (Figure S3A-D). Immunofluorescence of H3K27me3 and H3K9me3 in p53 KO cells showed a significant increase in Dac treated cells, compared to control, while Aza treated cells demonstrated an enrichment for H3K27me3 and a depletion of H3K9me3 signal. However, when compared to WT data, both treatments and the untreated control show a significant loss of both H3K27me3 and H3K9me3 staining, linking the absence of *p53* to a dramatic depletion on total H3K27me3 and H3K9me3 levels (Figure S3E, F).

**Figure 3:**
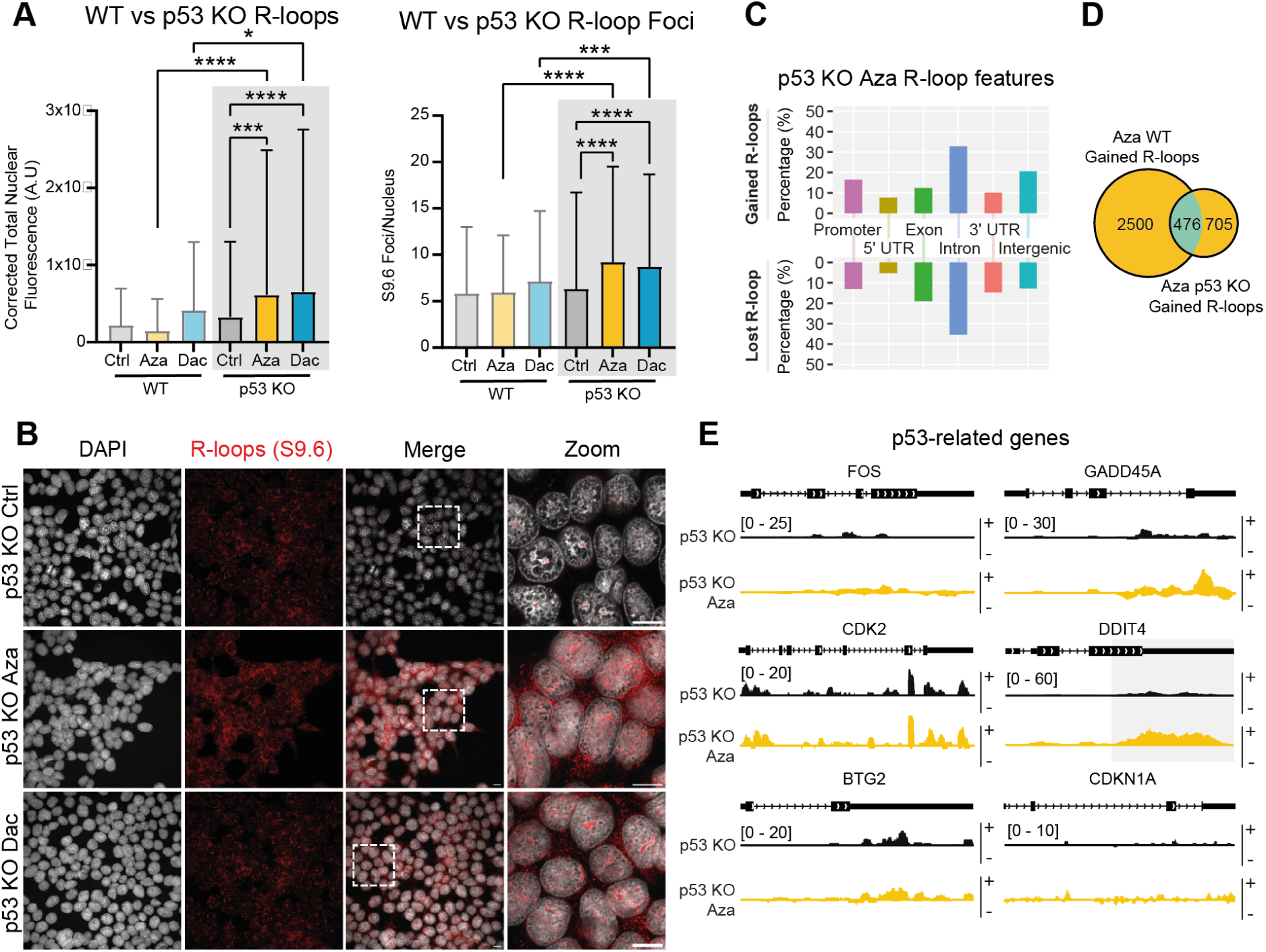
Removal of p53 re-models the epigenetic response to demethylating agents. (**A-B**): S9.6 staining in HEK293 p53 KO cells. (**A**) Analysis of R-loop S9.6 staining; corrected total nuclear fluorescence and S9.6 foci number per nucleus, in HEK293 p53 KO cells treated with 1.5μM Azacitidine (Aza), Decitabine (Dac) or solvent control for 48hrs (grey box denotes new data) vs the same treatment in HEK293 cells (WT) (data previously shown, see Figure 1). The p values were calculated by Kruskal-Wallis Test. *p < 0.05, ***p < 0.001, ****p < 0.0001. Error bars are 士SD. (**B**) Representative confocal images using S9.6 antibody to stain for R-loops (red) and DAPI as a counter stain for DNA (grey) in HEK293 p53 KO cells. Scale bars 10μm. (**C**) DRIPc-Seq feature distribution for significantly increased (gained) and decreased (lost) R-loop peak signals in cells treated with Aza. (**D**) Overlap of gained R-loop peaks in Aza-treated WT cells vs Aza-treated p53 KO cells. (**E**) DRIPc-Seq R-loop peak tracks of p53-related genes in p53 KO control-treated cells vs p53 KO Aza-treated cells, + and - symbols next to peak tracks denote the positive (+) and negative (−) (Watson & Crick) DNA strands.

Due to these substantially altered staining patterns in both R-loop and repressive chromatin markers, DRIPc-Seq was repeated in the HEK293 p53 KO cells, demonstrating a significant change in the R-loop profile of Aza treatment KO compared to WT. In p53 KO Aza treated cells, the highest percentage of R-loop gain sites were found in intronic regions (33.0%) with those gained in promoter regions only making up 13.7% (Figure 3C), in contrast to the 29% of newly gained promoter R-loops observed in the WT cells (Figure 2B). The enrichment of R-loops in the promoters of p53-target genes, which had been so evident in WT cells (Figure 2D, E), was lost in KO cells (Figure 3D, E). Our DRIPc-Seq analysis of Dac treated p53 KO cells encountered variability, which was sustained through both biological and technical replication (Figure S3G-J).

### R-loop interactome

Following the observed epigenetic landscape re-modelling after both the exposures of Aza and Dac, and from the absence of p53, we sought to determine the role R-loops were playing in these responses. We reasoned that a broad scale epigenetic re-mapping was occurring in response to these drugs, and that to achieve this, recruitment of the machinery that lays down and modifies epigenetic marks would have been required. Using DNA-RNA immunoprecipitation followed by Mass Spectrometry (DRIP-Mass), we assessed the proteins that were binding to R-loops in the differing conditions, to test this hypothesis (Figure 4A, S4A). The bulk of proteins were determined to be of nuclear origin (Figure 4B), however an increased prevalence of cytosolic proteins was observed in KO cells following Aza treatment (Figure 4B, S4A). Enrichment analysis of the WT response to Aza and Dac demonstrated an accumulation of epigenetic re-modelling proteins and complexes, including those components responsible for H3K9me3 and H3K27me3 writing (Figure 4B-D, S4A, B). Both conditions demonstrated accumulation of DNA methyltransferase proteins, while Dac was also found to accumulate histone acetylation writers (Figure 4D). In KO cells, while Dac exposure also induced the enrichment of epigenetic readers and writers, as had been seen in WT cells (Figure 4B-D, S4C), an overlapping response between Aza WT and KO cells was less evident, with no clear enrichment for the epigenetic remodelers in KO cells (Figure 4A-C, S4A-C). In fact, for Aza treated KO cells, this was the only condition where more total proteins interacting with R-loops had decreased from control conditions (Figure 4A, S4C).

**Figure 4:**
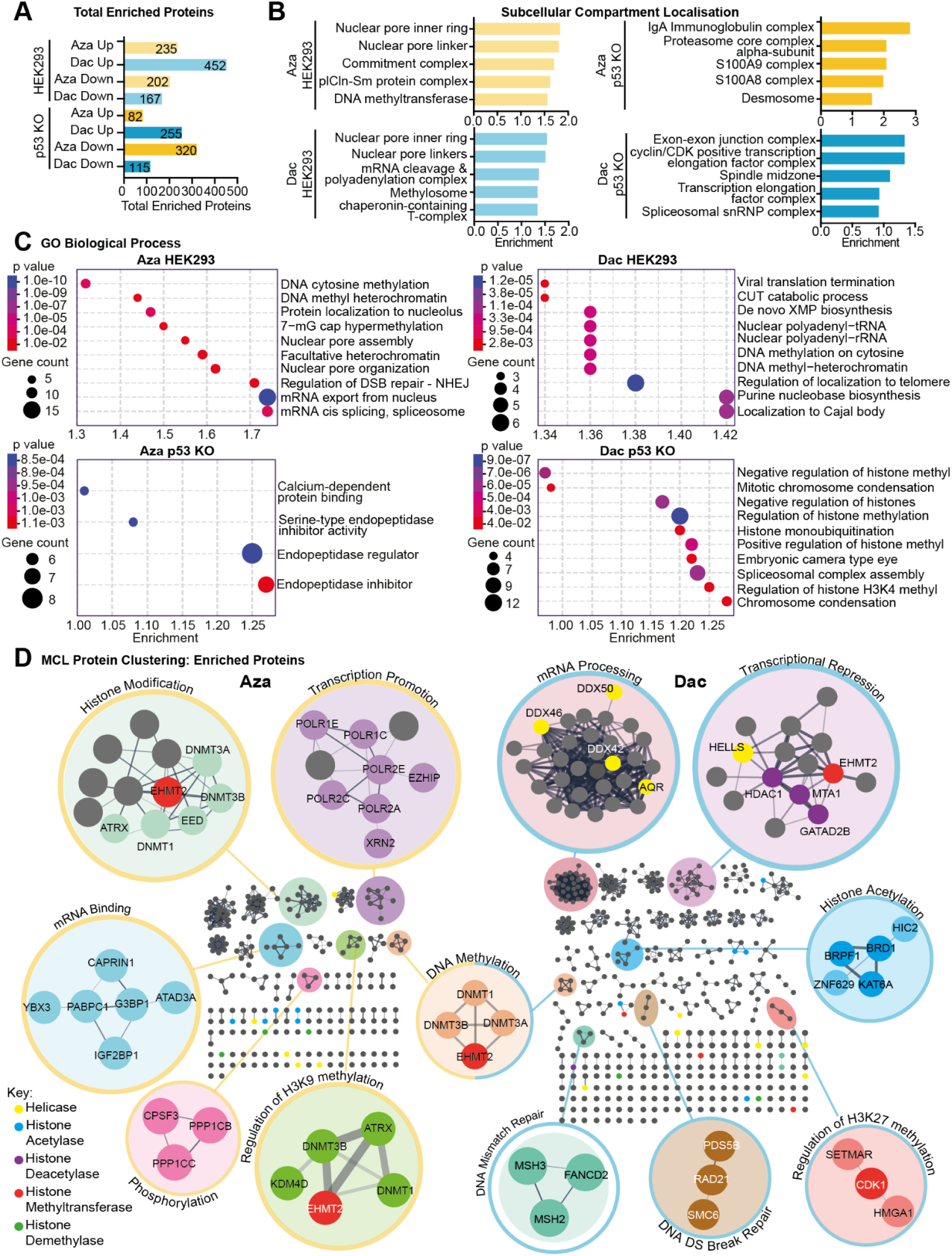
R-loops act as a recruitment site for epigenetic modifiers and damage response elements. **(A**) Total number of significantly enriched (up) or significantly depleted (down) proteins from Mass-Spectrometry of the R-loop interactome in HEK293 and HEK293 p53 KO cells treated with 1.5µM Azacitidine (Aza) or Decitabine (Dac), compared to solvent treated cells. (**B**) Subcellular compartment analysis of significantly enriched proteins. Bar plots show top five identified subcellular compartments by their Log10(observed/Expected) enrichment score. (**C**) Bubble plots of top ten hits from GO Biological term enrichment analysis of significantly enriched proteins. (**D**) STRING functional protein-protein interaction networks of significantly enriched proteins from Aza/Dac treated HEK293 cells, clustered using the Markov cluster algorithm (MCL) (38) via clusterMaker2 (72). Clusters and proteins of interest are highlighted using colours and annotated using their top biological function hit. Note that the same protein can appear in multiple clusters. Border colours denote clusters which are unique to Aza-treated cells (yellow) or Dac-treated cells (blue), an example of a shared cluster is shown in the centre (DNA Methylation).

### p53 rescue

It has recently been shown that loss of p53 reduces the level of S-adenosylmethionine (SAM), an important methyl-donor, resulting in a global loss of H3K9me3 (17). This places p53 as a powerful regulator of chromatin architecture and supports the previously shown loss of H3K9me3 and H3K27me3 observed in the HEK293 p53 KO cells (Figure S3E-F). In order to further solidify this relationship between p53 loss and re-modelling of chromatin markers, as well as explore a role for p53 in regulating R-loop formation, we utilised the ip53 H1299 cell line. This cell line possesses a *p53 Tet-on* inducible system allowing for restoration of p53 expression. In standard conditions, *p53* is transcriptionally silenced, but can be rescued through the addition of doxycycline (Dox) (Figure 5A, B) (35). As had been evident in the HEK293 cell line, for the ip53 H1299 cells in the absence of p53, R-loop staining intensity was observed to increase following Aza and Dac exposure (Figure 5C). Upon restoration of *p53,* the Aza and Dac induced accumulation of R-loop staining intensity was not observed, with no significant differences from either Dox^+^ or Dox^-^ control cells (Figure 5D-E), demonstrating a reversibility of this global accumulation of R-loop burden. Restricted H3K9me3 staining was detected in Dox^-^ Ctrl cells, which were further diminished following Aza and Dac exposure (Figure 5F). As had been seen for the R-loop staining, return of *p53* to control cells was found to rapidly restore both global H3K9me3 levels and a wild type like response to Aza and Dac exposure (FigureG-H, S3F). This effect was further observed in H3K27me3 staining, whereby rescue of p53 levels restored staining patterns and an expected response to both drugs (Figure 5I-K, S3E).

**Figure 5:**
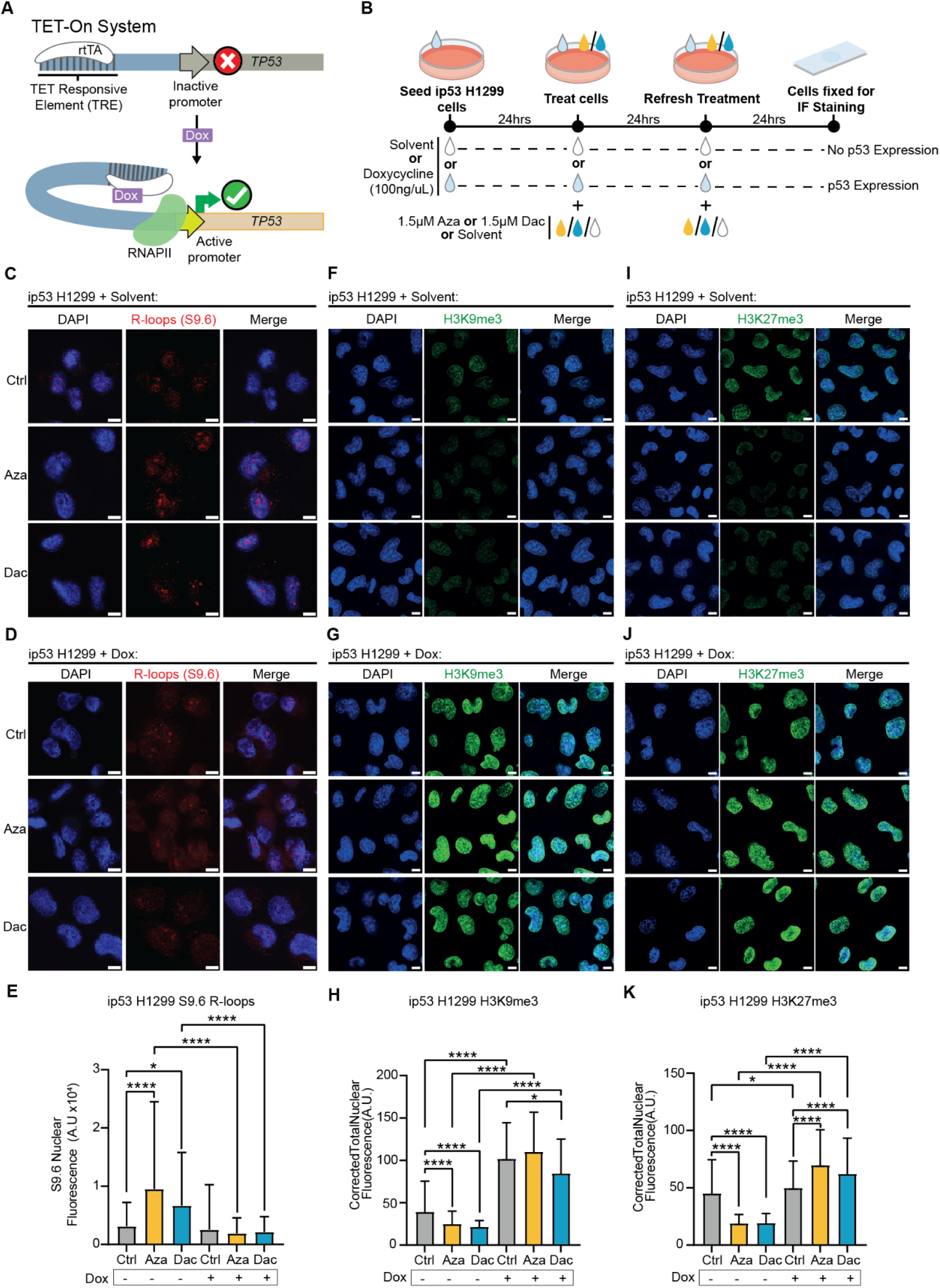
Restoration of p53 expression reverses epigenetic dysregulation and the response to demethylating agents. (**A**) Graphic of the TET-ON system for doxycycline induced p53 expression. In the presence of Doxycycline expression of the target gene (*TP53*) is reversibly activated. A Tet-responsive element (TRE) sits upstream of the *TP53* gene which contains an inactive promoter. The transcription factor rtTA recognises the TRE but can only induce gene expression in the presence of tetracyclines such as Doxycycline (Dox). When Dox is present it results in a conformational change allowing rtTA to recruit RNAPII, resulting in transcription of the *TP53* gene. (**B**) Graphic of the experimental design. ip53 H1299 cells were seeded and treated with solvent or 100ng/uL of Dox, to induce p53 expression, this treatment was refreshed every 24 hours. 24hrs after seeding, cells were treated with 1.5μM Azacitidine (Aza), Decitabine (Dac) or solvent, this treatment was then refreshed 24hrs later for a total of a 48hr treatment. Cells were then fixed and stained for immunofluorescence analysis. (**C-E**): Immunofluorescence staining of R-loops. (**C**-**D**) Representative confocal images using S9.6 antibody to stain for R-loops (red) and DAPI as a counter stain for DNA (blue) in ip53 H1299 cells. Cells treated with Dox express p53, whilst cells treated with solvent do not. Scale bars 10μm. (**E**) Analysis of R-loop S9.6 staining; corrected total nuclear fluorescence in ip53 H1299 cells treated with or without Dox and either 1.5μM Aza, Dac or solvent control for 48hrs. The p values were calculated by Kruskal-Wallis Test. *p < 0.05, ***p < 0.001, ****p < 0.0001. Error bars are 士SD. (**F-H**) Immunofluorescence staining of H3K9me3. (**F-G**) Representative confocal images of H3K9me3 staining (green) and DAPI as a counter stain for DNA (blue) in ip53 H1299 cells. Cells treated with Dox express p53, whilst cells treated with solvent do not. Scale bars 10μm. (**H**) Analysis of H3K9me3 staining; corrected total nuclear fluorescence in ip53 H1299 cells treated with or without Dox and either 1.5μM Aza, Dac or solvent control for 48hrs. The p values were calculated by Kruskal-Wallis Test. *p < 0.05, ***p < 0.001, ****p < 0.0001. Error bars are 士SD. (**I-K**) Immunofluorescence staining of H3K27me3. (**I-J**) Representative confocal images of H3K27me3 staining (green) and DAPI as a counter stain for DNA (blue) in ip53 H1299 cells. Cells treated with Dox express p53, whilst cells treated with solvent do not. Scale bars 10μm. (**K**) Analysis of H3K27me3 staining; corrected total nuclear fluorescence in ip53 H1299 cells treated with or without Dox and either 1.5μM Aza, Dac or solvent control for 48hrs. The p values were calculated by Kruskal-Wallis Test. *p < 0.05, ***p < 0.001, ****p < 0.0001. Error bars are 士SD.

## Discussion

Following observations that both DNA and RNA methylation content can influence the propensity to form R-loops, we chose to evaluate the chemical removal of these marks, and whether these differentially impacted upon R-loop patterns across the genome. Our finding that p53 status is a critical factor in the response of cells to both Aza and Dac (Figure 5C-E, 6) was unexpected, as p53 activation for either of these drugs had not been directly reported, while a link between R-loop modulation and p53 had not been demonstrated. The response to Aza was dependent on the activation of p53 (Figure 2F), including the highly enriched appearance of R-loops in the promoters of p53-target genes (Figure 2D-E). Such R-loops were not accompanied by detectable changes to expression of these loci, while for the 111 differentially expressed genes that were detected following Aza treatment, only 3 were found to be enriched for an R-loop (Figure S2F, H). The cellular response in the case of Aza treatment was sufficient to protect against the damaging impact of the drugs on genomic stability, which was lost upon removal of p53 (Figure 3C-E, S3A-D). Interestingly, in the absence of p53, a larger proportion of associating proteins following Aza treatment were found to be cytosolic, rather than the other conditions where nuclear proteins were highly enriched (Figure S4B, C). Novel work has recently identified the presence of cytosolic R-loops in specific conditions (54), with these aberrant structures associated with cell death. Further work would be required to determine if Aza was inducing an accumulation of these cytosolic R-loops. In contrast, the response to Dac was not determined through activation of p53 (Figure 2F), but by the initial state of heterochromatin and then subsequently from an ability to remodel repressive histone modifications following exposure (Figure S3E-F, 4C-D, 5F-K). The fact we see only a handful of significant transcriptional changes following Dac treatment (Figure 1E) is testament to this re-modelling, and the ability of the epigenome to adapt and respond to stimuli, further demonstrating the multi-layered epigenetic control that is employed on gene expression. Our analysis of the proteome that was associated with these regions of the genome demonstrated a clear link between R-loops and the re-modelling of global histone marks. Following drug exposure, recruitment of these repressive histone modifications was required to mitigate against the accumulative damaging impact. In the case of Dac, such recruitment was insufficient to protect the cell from DNA damage, as exemplified by heightened micronuclei formation and errors in mitosis (Figure S3A-D). When the initial heterochromatin state was destabilized by the removal of p53 (Figure S3E, F), this damage was amplified for Dac treatment, while also seen in Aza treated samples (Figure S3A-D).

While we have attempted to understand the global epigenetic consequences of exposure to these demethylating agents, the direct genetic function of much of the changes to the global R-loop distribution remains opaque. We found a clear gain of promoter R-loops following Aza exposure, while terminator R-loops were especially enriched in Dac treated samples (Figure 2B, C). Promoter R-loops are generally thought to increase transcription (23) while terminator R-loops are known to often be transcriptionally repressive and frequently damage inducing, a feature that was detected in Dac treated cells (Figure S1D-G) (28, 29). Globally, we did not detect a clear relationship between the gain of R-loops and the location of newly demethylated sites, for either drug (Figure S2E). This of course begs the question on the functionality of these R-loops, especially given the previous observation that transcriptionally active un-methylated CpG islands are enriched for R-loops (21). While our analysis cannot preclude that genome-wide changes to splicing patterns will have occurred, we did determine that the changes in transcription that were detected, were not accompanied by a newly formed R-loop at that locus (Figure S2F). We speculate that these newly formed R-loops represent a novel component of the epigenetic response and re-modelling that the cell undergoes following the aberrant removal of nucleic acid methylation. This provides a recruitment point for the machinery that lays down repressive histone modifications and DNA damage response elements (Figure 4B-D), which together serve to maintain the appropriate transcriptional dosage and epigenetic architecture of critical loci in the genome.

A limitation of our study is that we were not able to completely characterize the differential sensitivity to these drugs, including interrogation of changes to RNA m5C levels, which would be expected to be inhibited by Aza, but not Dac. Sequencing of this mark has been historically technically challenging and the most widely used sequencing approach in mammalian systems involves the treatment of samples with Aza, precluding its applicability to our experiments (51, 55). Recent works have highlighted that RNA modifications can preferentially dispose the modified RNA to form an R-loop (56, 57), and therefore we might hypothesize that the impaired propensity to form specific R-loops following global removal of RNA m5C may instigate the downstream epigenetic responses that we observed following Aza treatment. Future studies will be required to explore this further. Separately, we highlighted the variability that was observed in DRIPc-Seq and the requirement for appropriate controls (Figure S2A). This variability was most profound in Dac treated conditions, and was further compounded in p53 KO settings, likely indicative of a biological effect (Figure 1E, S3G-J). However, previous work has highlighted that the S9.6 antibody can be subject to technical challenges (49, 58), and our findings provide further evidence that controlled use of this antibody is imperative, with RNAse H controls and caution taken when reading across experimental replicates.

To the best of our knowledge, p53 status is not currently considered clinically when determining treatment with Aza or Dac in AML or MDS (59, 60). Conflicting data exists as to the impact of loss of p53 and the response to Aza and Dac treatment, however there is some consensus that loss of p53 may be an indicator of resistance to Aza treatment, while Dac induced apoptosis is thought to be enhanced by the loss of p53 (61–70). Interestingly, substantial effort is currently being employed in utilising Aza in particular, in combination therapies, including cases of MDS which are not responsive to demethylating agents in isolation. A clinical trial has demonstrated the substantially improved efficacy of Aza when used in combination with Eprenetapopt (APR-246), a first in class small molecule that can restore wt p53 function in null settings (71). Our data presented here supports these observations and provides mechanistic insights as to the underlying causes (Figure 6), linking the Aza-specific activation of p53, and subsequent re-modelling of the epigenome to the downstream apoptotic consequences. Perhaps our most surprising finding was the reversibility of the effects seen following p53 removal, both in the architecture of the epigenome and the response to the drugs, which further aligns with the observed efficacy with Eprenetapopt. The staining distribution of H3K9me3 and H3K27me3 was substantially ablated in the two independent p53 null cell lines, while R-loop staining was enriched. Restoration of p53 was sufficient to rapidly reverse the destabilisation of the epigenetic landscape and restore the burden of R-loops following Aza and Dac treatment to a wild type-like response. Given the significant breadth of epigenetic therapies currently being trialled for various disease states, the dynamism displayed in the epigenome displayed here, that enables reversibility in ways that are not possible with genetic changes, should further encourage this area of research.

**Figure 6:**
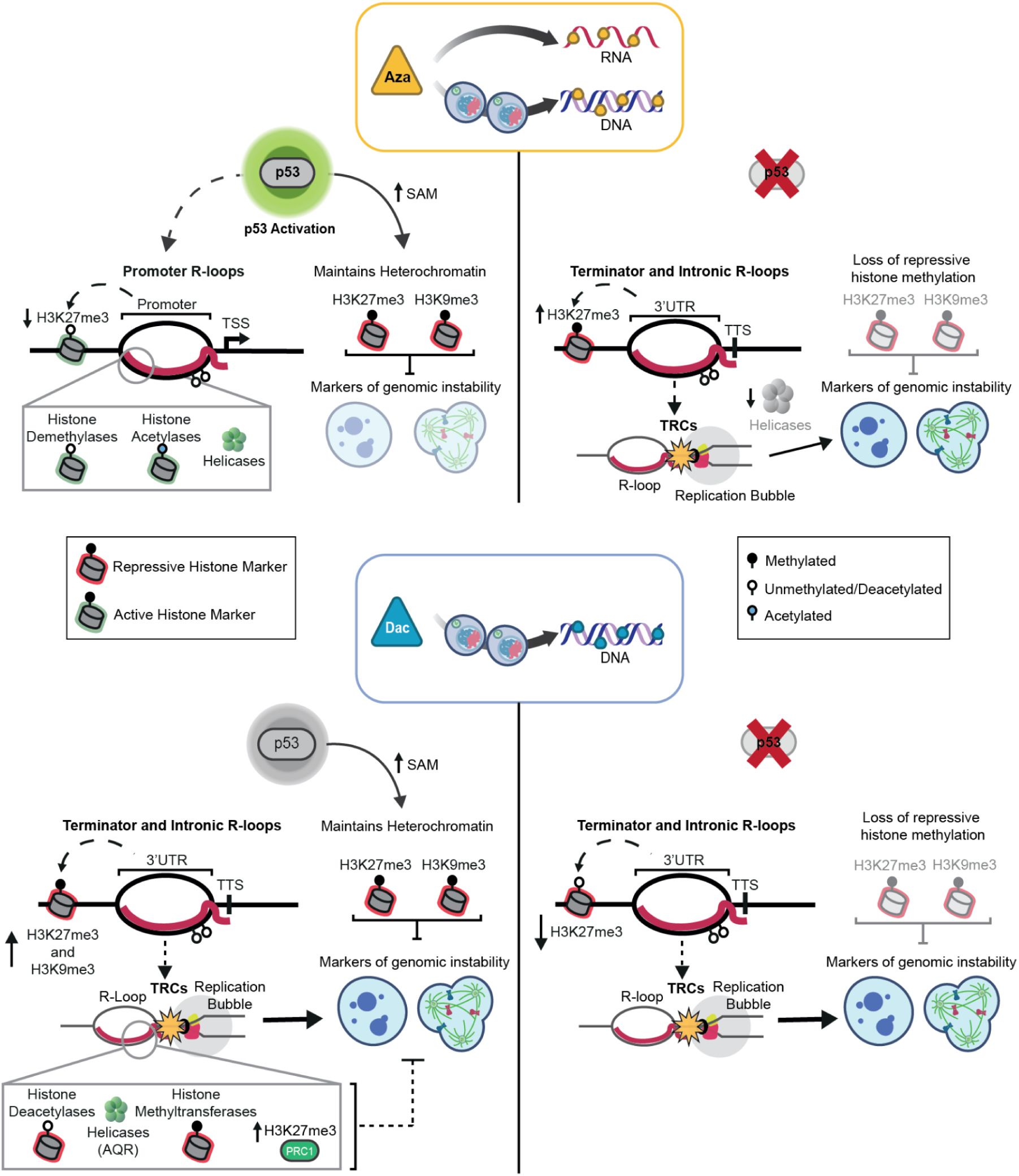
Summary of observed Aza and Dac phenotypes in WT and p53 KO cells. Top: In WT cells (left side), Azacitidine (Aza) treatment resulted in an increase in active p53. There was also an observed enrichment of R-loops across gene promoters alongside changes in histone methylation, a significant decrease in H3K27me3 and the recruitment of histone modifiers and helicases. There was no observed significant increase in micronuclei or aberrant mitosis. It has been shown that p53 is important for maintaining levels of S-adenosylmethionine (SAM) which was shown to be linked to a global reduction in H3K9me3 (17). Repressive histone markers H3K9me3 and H3K27me3 are also known to be important in preventing genomic instability (73). In p53 KO cells (right side), Aza treatment resulted in an enrichment of R-loops at the 3’UTR and across introns, alongside a significant increase in H3K27me3. We hypothesise that these R-loops are involved in transcription-replication conflicts (TRCs) which is driving the observed significant increase in markers of genomic instability. There was also a reduction in the recruitment of helicases to R-loops. p53 KO cells were also observed as having a significant reduction in repressive histone markers. Bottom: In WT cells (left side), Decitabine (Dac) did not significantly increase levels of active p53. Dac treatment resulted in an enrichment of R-loops across gene terminators and introns, an increase in repressive histone markers and an increase in markers of genomic instability. We hypothesise that the enrichment of these R-loops results in increased TRCs which is driving genomic instability. There was observed enriched recruitment of histone modifiers, helicases such as Aquarius (AQR), which is associated with DNA repair (74), and members of the Polycomb Repressive Complex 1 (PRC1) which is important for maintaining H3K27me3 (75). In p53 KO cells (right side), Dac-treatment resulted in enrichment of R-loops at gene terminators and introns, a significant decrease in H3K27me3 and H3K9me3, a significant increase in markers of genomic instability. There was also a decrease in recruitment of histone modifiers and helicases.

## Authors Contributions

EH and MVDP conceptualised the study and wrote the manuscript. EH performed the experiments with support from GO, SS and RD. AW analysed the data generated. All authors have read and approved the manuscript.

## Acknowledgements

We acknowledge Ivano Amelio (Universität Konstanz) for helpful discussions in the planning phase. We acknowledge the support of Lajos Kalmar, Lucia Pinon and Catarina Franco (MRC Toxicology Unit) for their support with data analysis, microscopy and mass spec experiments respectively. Figure 6 was generated with Biorender.com using an Academic Licence.

---

**Supplementary Figure 1.**
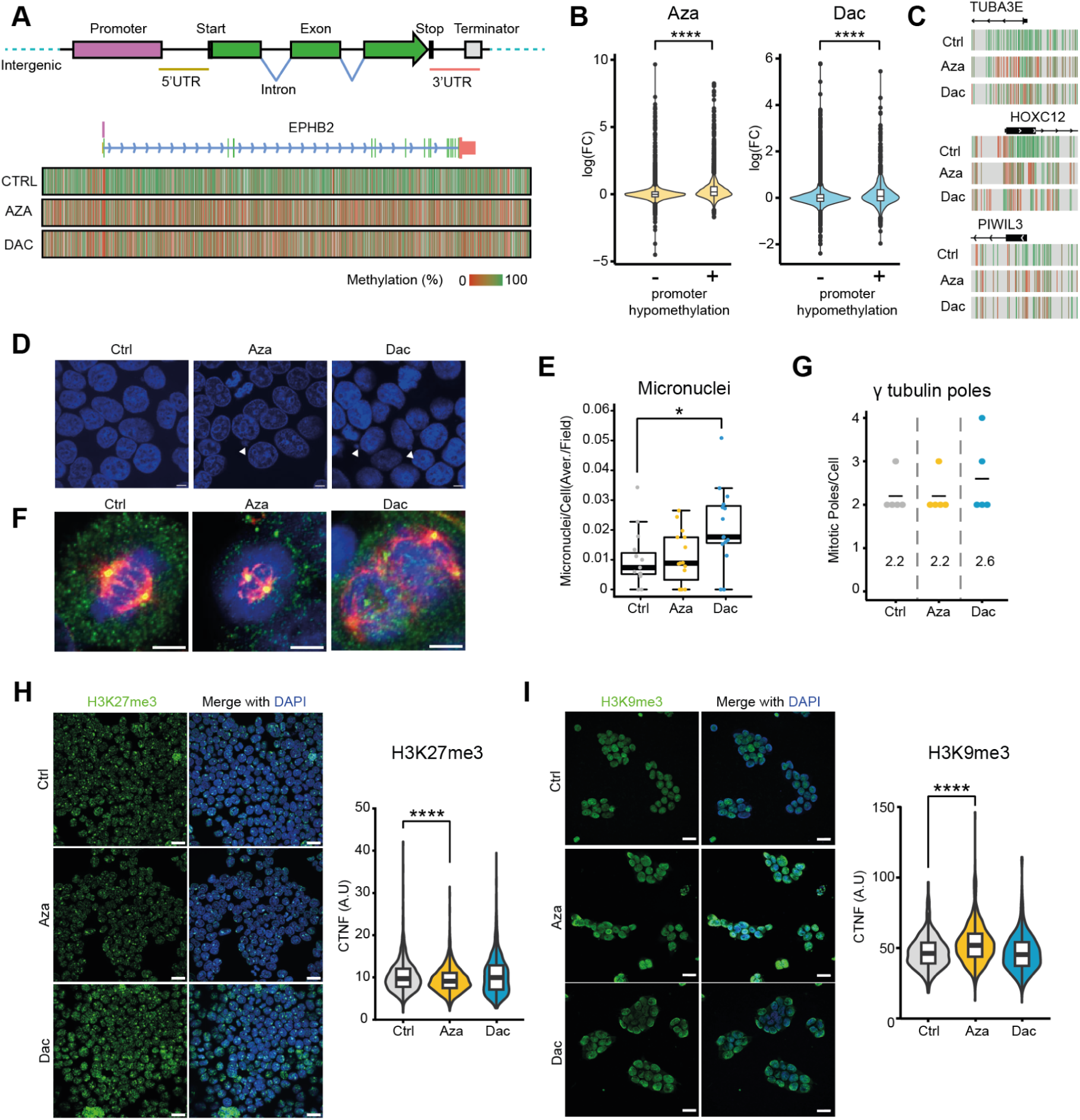
(**A**) Schematic cartoon of gene architecture and a heatmap representation of changes in CpG methylation upon Azacitidine (Aza) and Decitabine (Dac) treatment of an exemplary gene. (**B**) Violin plot of RNA-seq log(FC) values of genes with (+) and without hypomethylated promoter (−) for Aza and Dac. (**C**) Examples of gene promoter regions hypomethylated following Dac and Aza treatment. (**D**-**E**): Micronuclei analysis. (**D**) Representative confocal images of micronuclei in HEK293 cells following treatment with 1.5μM Aza, Dac or solvent control. White arrows indicate location of micronuclei, scale bar 10μm. (**E**) Box plot shows micronuclear quantification (count per cell), each dot represents a field. The p values were calculated by one-way ANOVA *p < 0.05, data are presented as mean 土SEM. (**F-G**): Mitotic pole analysis. (**F**) Representative confocal images of α-tubulin (red) and γ-tubulin (green) staining in HEK293 cells treated with 1.5μM Aza, Dac or solvent control. Scale bar 10μm. DAPI (blue) used to counterstain DNA. (**G**) Dot plots displaying quantification of mitotic poles; each dot represents a mitotic cell. Black line indicates the mean. (**H**): H3K27me3 analysis. Representative confocal images of H3K27me3 staining (green) in HEK293 cells following treatment with 1.5μM Aza, Dac or solvent control. DAPI used to counter stain DNA (blue), scale bar 10μm. Violin plot of H3K27me3 corrected total nuclear fluorescence (CTNF). Statistical testing was calculated by Kruskal-Wallis Test, ****p < 0.0001. (**I**): H3K9me3 analysis. Representative confocal images of H3K9me3 staining (green) in HEK293 cells following treatment with 1.5μM Aza, Dac or solvent control. DAPI used to counter stain DNA (blue), scale bar 10μm. Violin plot of H3K9me3 CTNF. The p values were calculated by Kruskal-Wallis Test, ****p < 0.0001.

**Supplementary Figure 2.**
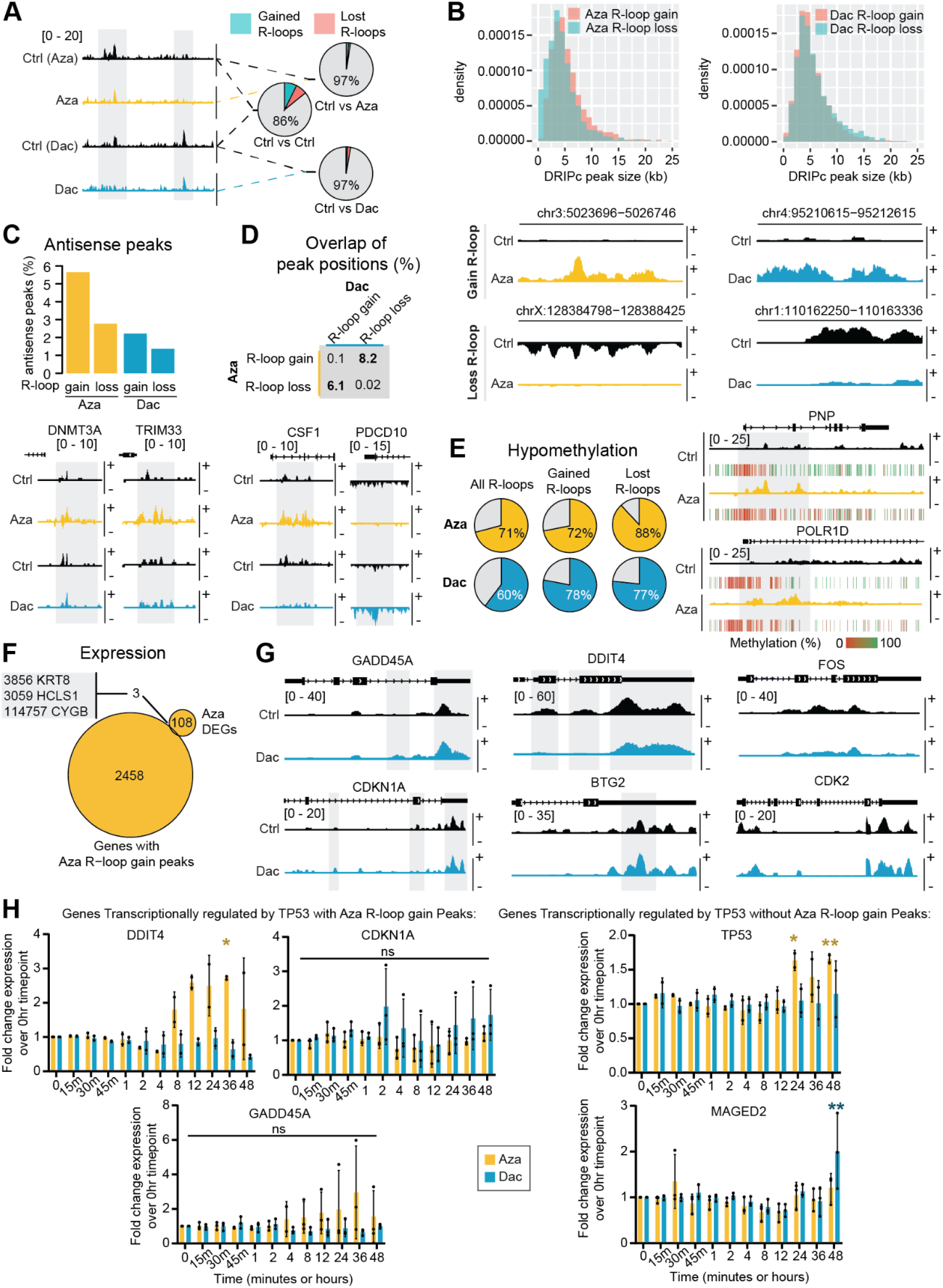
(**A**) Examples of DRIPc-Seq R-loop peak tracks for Azacitidine (Aza) and Decitabine (Dac) treated HEK293 cells, alongside each treatment’s corresponding solvent control treated group. Pie charts of the proportion of significantly changed R-loop peaks (gained, lost and no change (grey)), between the two control treatments and Aza/Dac treatment with their respective control treatment. (**B**) Size distribution of significantly changed DRIPc-peaks (gained/lost) for Aza and Dac treatment. Gene plots below are examples of a gained and a lost R-loop signal in Aza and Dac-treated cells compared to control-treated cells. (**C**) Proportion of antisense peaks of all significantly changed DRIPc-seq R-loop peaks (gained/lost) under Aza and Dac treatment with two examples of Aza-exclusive promoter-region antisense peak formation shown below (antisense strand is denoted by the -, sense strand is +). (**D**) Overlap matrix between Aza and Dac R-loop gain and loss peak positions with two examples of regions with an Aza R-loop gain and Dac R-loop loss overlap and a Dac-R-loop gain and Aza-R-loop loss overlap shown below. (Watson/forward-strand R-loop (+), Crick/reverse-strand R-loop (−)) (**E**) Proportion of hypomethylation of genetic regions of all R-loop peaks and for differentially changed R-loop peaks (gain/loss) under Aza and Dac treatment with two examples of Aza R-loop formation in an unmethylated promoter region. (**F**) Venn diagram of genes which contain Aza-induced gained R-loop formation and differentially expressed up-regulated genes following Aza treatment. (**G**) DRIPc-seq peak data for representative p53-related genes after Ctrl and Dac treatment. (**H**) Fold change gene expression of five p53-target genes, identified as being with or without gained Aza-induced R-loops, over time in Aza or Dac treated HEK293 cells, compared to untreated cells (0 time point). n = three biological repeats. The p values were calculated by one-way ANOVA, *p < 0.05. **p < 0.01. Error bars are 士SEM.

**Supplementary Figure 3.**
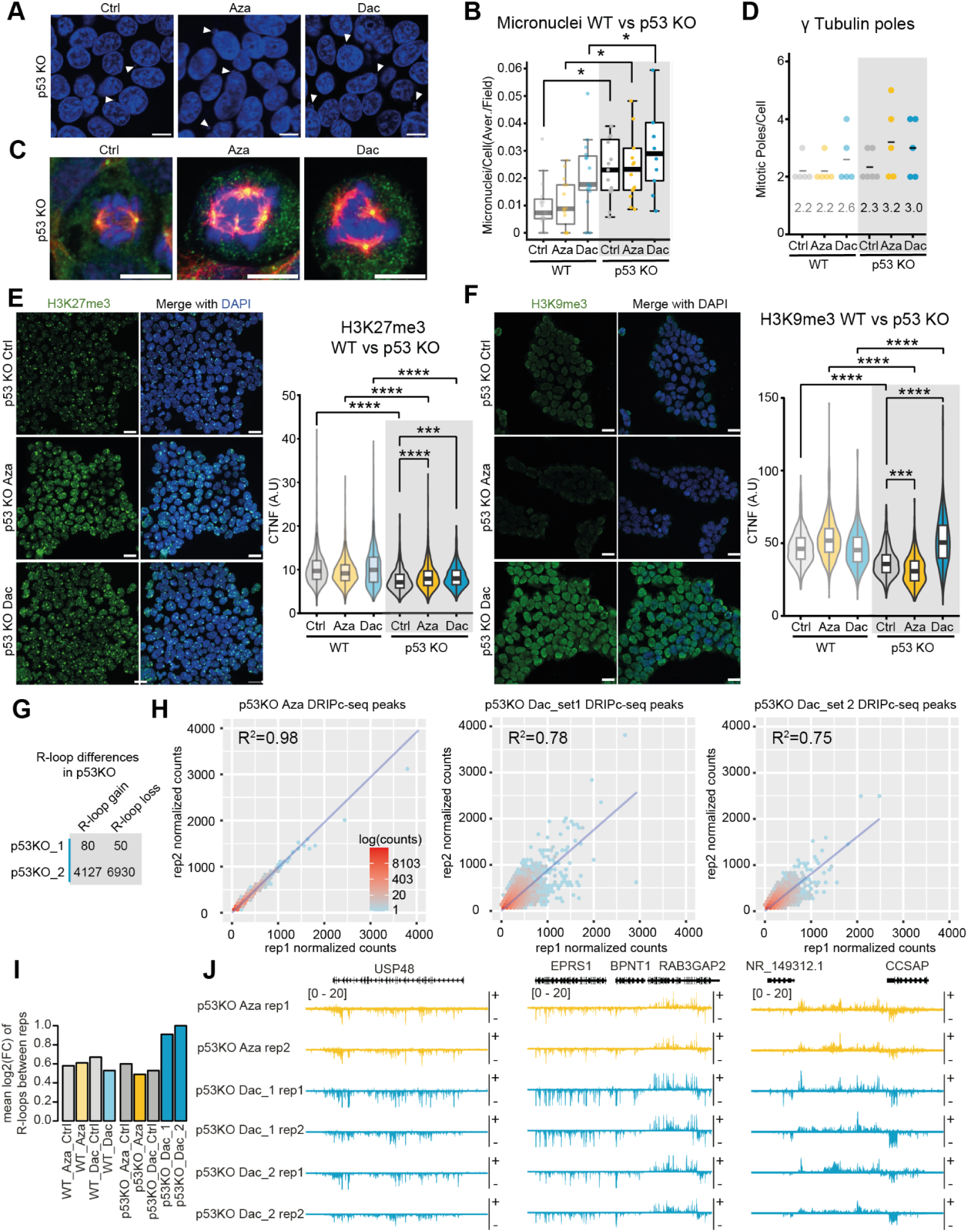
(**A-B**): Micronuclei analysis. (**A**) Representative confocal images of micronuclei in HEK293 p53 KO cells following treatment with 1.5μM Azacitidine (Aza), Decitabine (Dac) or solvent control. White arrows indicate the location of micronuclei, scale bar 10μm. (**B**) Box plot shows micronuclear quantification (count per cell), in HEK293 (WT) cells (data previously shown, see Figure S1) vs the same treatment in HEK293 p53 KO cells, grey box denotes new data, each dot represents a field. The p values were calculated by one-way ANOVA with Tukey’s correction. *p < 0.05, data are presented as mean 土SEM. (**C**-**D**): Mitotic pole analysis. (**C**) Representative confocal images of α-tubulin (red) and γ-tubulin (green, when overlapping with α-tubulin stain appears yellow) staining in HEK293 p53 KO cells treated with 1.5μM Aza, Dac or solvent control. Scale bar 10μm. DAPI (blue) is used to counterstain DNA. (**D**) Dot plots displaying quantification of mitotic poles in HEK293 cells (data previously shown, see Figure S1) vs HEK293 p53 KO cells, grey box denotes new data, each dot represents a mitotic cell. Black line indicates the mean. (**E**) H3K27me3 analysis. Representative confocal images of H3K27me3 staining (green) in HEK293 p53 KO cells following treatment with 1.5μM Aza, Dac or solvent control. DAPI is used to counter-stain DNA (blue), scale bar 10μm. Violin plot of H3K27me3 corrected total nuclear fluorescence (CTNF), in HEK293 cells (data previously shown, see Figure S1) vs HEK293 p53 KO cells, grey box denotes new data. The p values were calculated by Kruskal-Wallis Test, ***p < 0.001, ****p < 0.0001. (**F**) H3K9me3 analysis. Representative confocal images of H3K9me3 staining (green) in HEK293 p53 KO cells following treatment with 1.5μM Aza, Dac or solvent control. DAPI is used to counter-stain DNA (blue), scale bar 10μm. Violin plot of H3K9me3 CTNF, in HEK293 cells (data previously shown, see Figure S1) vs HEK293 p53 KO cells, grey box denotes new data. The p values were calculated by Kruskal-Wallis Test, ***p < 0.001, ****p < 0.0001. (**G**) Table to compare the number of significantly increased R-loop peaks (gain) and significantly decreased R-loop peaks (loss) identified by DRIPc-Seq in control-treated HEK293 p53 KO samples 1 and 2. (**H**) XY correlation heatmaps of DRIPc-Seq signal in replicate 1 vs replicate 2 of Aza and Dac-treated HEK293 p53 KO cells with R2 linear regression as a measure of variation between replicates. (**I**) Mean log2 (fold change) of R-loop signals between sample repeats of all DRIPc-Seq samples included in this study. (**J**) Example gene regions of DRIPc-Seq R-loop peak tracks of both replicates for Aza and Dac-treated HEK293 p53 KO cells, + and - symbols next to peak tracks denote the positive (+) and negative (−) (Watson & Crick) DNA strands.

**Supplemental Figure 4.**
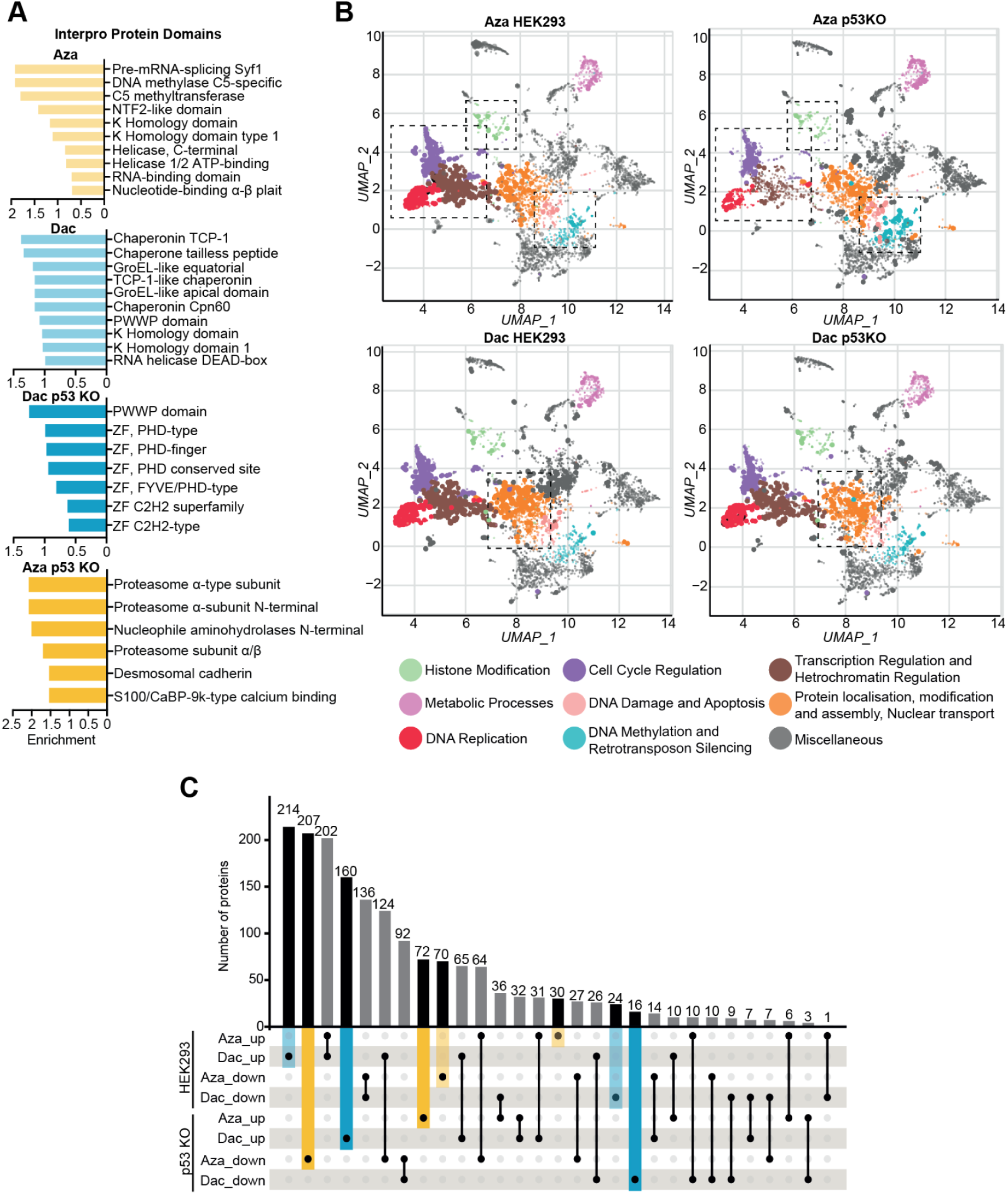
(**A**) InterPro protein domain analysis of significantly enriched proteins from Mass-spectrometry of the R-loop interactome in HEK293 and HEK293 p53 KO cells treated with 1.5µM Azacitidine (Aza) or Decitabine (Dac), compared to solvent treated cells. (**B**) Scatter plots of DRIP-Mass Spectrometry proteins from Aza or Dac treated HEK293 and HEK293 p53 KO cells, clustered based on GO biological enrichment term analysis. Clusters are computed using the Leiden algorithm, with points plotted on the first two UMAP dimensions. Larger and more opaque points represent more significantly enriched terms. Clusters of interest are highlighted with colours, all other clusters are coloured grey. Some clusters are further highlighted by dashed boxes to allow for easy comparison between the drug conditions (**C**) Upset-Plot showing the number of unique (black/coloured bars) and shared (grey) significantly enriched (up) and significantly depleted (down) proteins from Mass-spectrometry of the R-loop interactome in HEK293 and HEK293 p53 KO cells treated with 1.5µM Aza or Dac.

**Supplementary Table 1:**
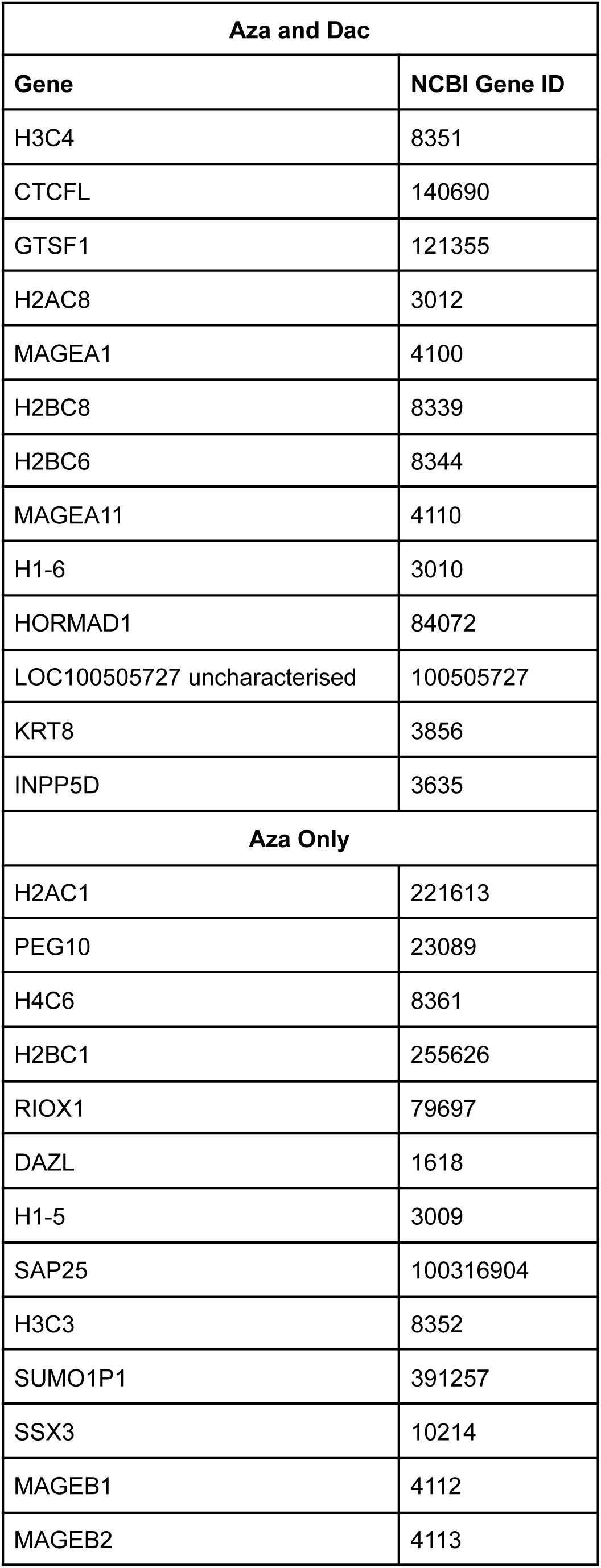
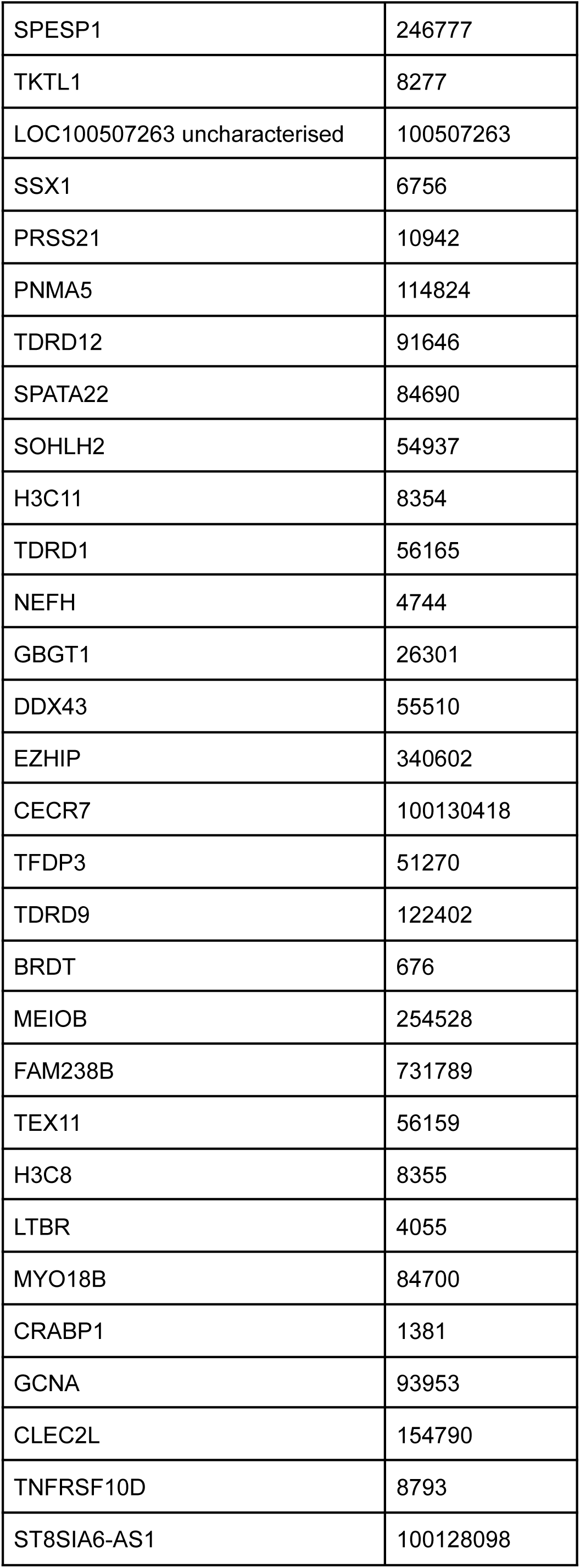
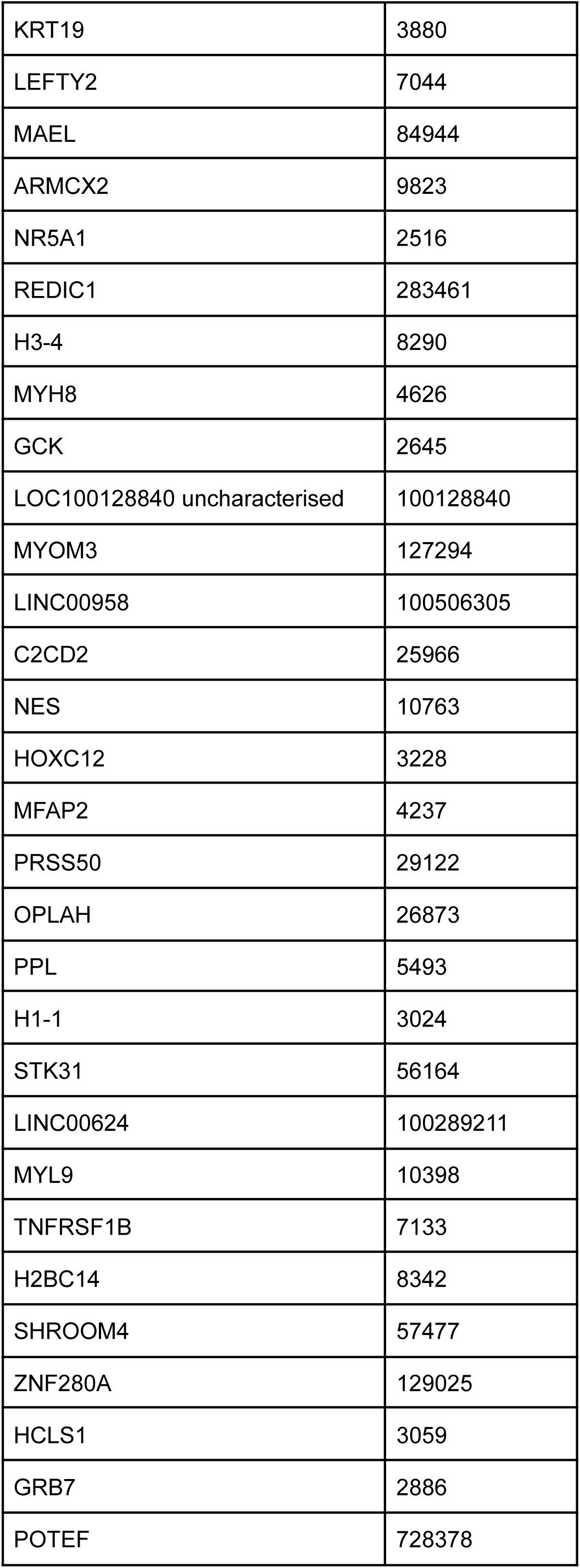
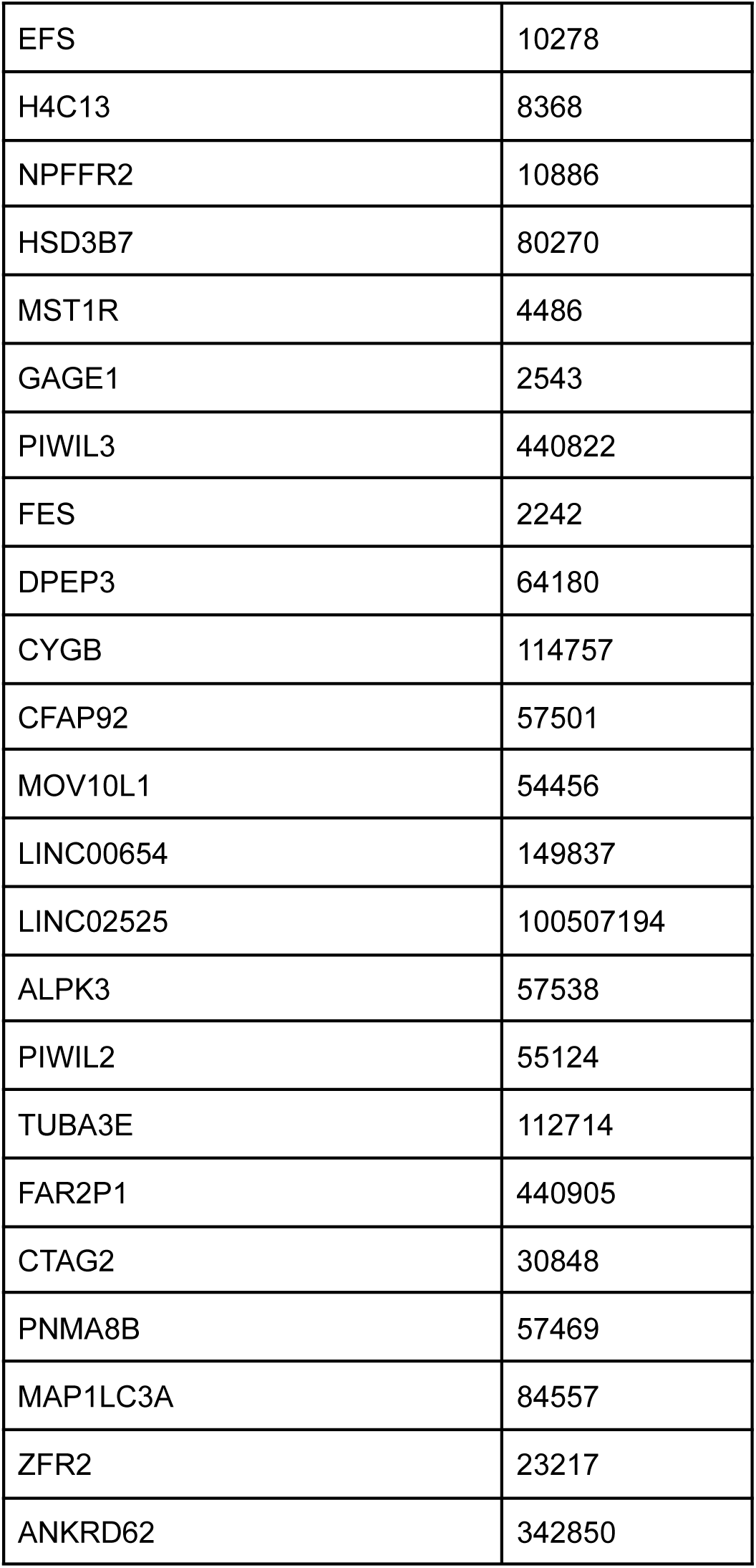
List of RNA-Seq Differentially Expressed Genes.

